# SARS-CoV-2 NSP14 governs mutational instability and assists in making new SARS-CoV-2 variants

**DOI:** 10.1101/2023.09.28.559966

**Authors:** Sk. Sarif Hassan, Tanishta Bhattacharya, Debaleena Nawn, Ishana Jha, Pallab Basu, Elrashdy M. Redwan, Kenneth Lundstrom, Debmalya Barh, Bruno Silva Andrade, Murtaza M. Tambuwala, Alaa A. Aljabali, Altijana Hromić-Jahjefendić, Wagner Baetas-da-Cruz, Vladimir N. Uversky

## Abstract

Severe acute respiratory syndrome coronavirus 2 (SARS-CoV-2), the rapidly evolving RNA virus behind the COVID-19 pandemic, has spawned numerous variants since its 2019 emergence. The multifunctional NSP14 enzyme, possessing exonuclease and mRNA capping capabilities, serves as a key player. Notably, single and co-occurring mutations within NSP14 significantly influence replication fidelity and drive variant diversification. This study comprehensively examines 120 co-mutations, 68 unique mutations, and 160 conserved residues across NSP14 homologs, shedding light on their implications for phylogenetic patterns, pathogenicity, and residue interactions. Quantitative physicochemical analysis categorizes 3953 NSP14 variants into three clusters, revealing genetic diversity. This research underscores the dynamic nature of SARS-CoV-2 evolution, primarily governed by NSP14 mutations. Understanding these genetic dynamics provides valuable insights for therapeutic and vaccine development.

## 1. Introduction

The emergence of SARS-CoV-2 in December 2019 had a profound global impact. This virus, a member of the Coronaviridae family, is a betacoronavirus characterized by its positive-sense, single-stranded RNA genome [1, 2]. Sequence analysis revealed striking similarities between SARS-CoV-2 and its predecessors, MERS-CoV (2013) and SARS-CoV (2001) [3]. Upon infection, SARS-CoV-2 can manifest with a wide spectrum of symptoms, ranging from pneumonia to severe acute respiratory syndrome [4]. Its genome encodes four structural proteins—Spike (S), Envelope (E), Membrane (M), and Nucleocapsid (N)-and sixteen non-structural proteins (NSP1 to NSP16), primarily involved in replication and protease activity [5, 6]. Due to its high transmissibility and devastating global impact, extensive research on the SARS-CoV-2 genome has led to the creation of valuable databases such as GISAID and NextStrain, providing crucial insights into its genetic makeup [7, 8, 9].

The SARS-CoV-2 genome, roughly 30 Kb in length, is dominated by two polyproteins, pp1a and pp1ab, which are subsequently cleaved into non-structural proteins. Among these, RNA-dependent RNA polymerase (RdRp) is central to RNA synthesis, forming a core complex with nsp7, nsp8, and nsp12, while nsp14 interacts with this complex, wielding essential proofreading activity [6, 10]. NSP14, consisting of approximately 527 amino acids, possesses a bifunctional role. Its N-terminal region harbors a 3’-5’ exonuclease responsible for proofreading during replication, ensuring genomic fidelity and reducing error rates [11, 12, 13]. The C-terminal N7-methyltransferase domain plays a critical role in mRNA capping and evading host immune responses [14]. Mutations in the DxG domain within this region can abolish RNA capping activity without affecting exonuclease function [15]. While RdRp facilitates viral replication, post-transcriptional processes involve capping and polyadenylation to protect the genome from degradation, evade antivirals, and enable efficient host translation. This is orchestrated by non-structural proteins NSP13, NSP14, and NSP16, with NSP14 typically associating with its accessory protein, NSP10 [16, 17].

The NSP10/14 complex plays a pivotal role in maintaining replication fidelity, controlling mutation rates, and influencing resistance to antiviral drugs [18]. Recent research highlights compounds like SGC0946 and SGC8158 that selectively target SARS-CoV-2 NSP14 by blocking SAM and RNA-binding sites [19].

RNA viruses, known for their high mutation rates, create diverse viral strains. These mutations can either enhance virulence, enable drug resistance, or go unnoticed without impacting transmission. The presence of a proofreading mechanism, like that found in SARS-CoV-2, can reduce mutation rates compared to other viruses, estimated at approximately 10^*−*^6 mutations per site/cycle [1]. Mutations in NSP14, particularly in the ExoN domain, contribute to increased genome-wide mutational loads [9]. Specific mutations, such as the shift from cytosine to uracil, can affect host inflammatory cytokine production [20]. Mutations in the active sites and zinc finger motifs of the Exonuclease domain can lead to lethal phenotypes [21, 11]. The significance of understanding these mutations lies in their impact on diagnostic assays, vaccine development, and viral transmissibility [22, 23]. Targeting NSP14, which maintains genomic stability, offers potential therapeutic strategies against SARS-CoV-2 [24]. Beyond its role in proofreading, NSP14 acts as a translation inhibitor in host cells, a characteristic conserved across human coronaviruses [23]. Recent studies have identified natural compounds that can target several key residues within NSP14 as potential targets for therapeutic interventions [25]. Conservation across coronaviruses is evident, with the NSP14 of SARS-CoV-2 sharing high amino acid similarity with its counterparts in SARS-CoV and MERS-CoV [23]. The ExoN and N7-MTase domains play vital roles in inhibiting host antiviral responses, and mutations in these domains can trigger host antiviral responses [23].

This study explores unique mutations in NSP14 across 13 different geographic locations and assesses their impact, differentiating between neutral and deleterious mutations using the PredSNP server [26].

## 2. Data Specifications and Methods

### 2.1 Data

A total of 1,182,629 SARS-CoV-2 NSP14 sequences, devoid of any ambiguous characters, were sourced from the GISAID-Virusurf database. Remarkably, among this extensive dataset, only 3,953 sequences, equivalent to a mere 0.33%, exhibited unique and distinct variations, representing the global diversity of SARS-CoV-2 NSP14. For reference, the NSP14 sequence denoted as YP 009725309 was retrieved from the NCBI.

### 2.2 Methods

#### 2.2.1 Analyze sequence variation

Single mutations in all the 3953 unique NSP14 sequences were determined using the Virus Pathogen Resource ViPR by inputting Fasta file of NSP14 sequences [27].

Furthermore, the predicted effect on pathogenicity of all the mutations was analyzed with PredictSNP and PhD-SNP [26, 28]. Note that PredictSNP web server makes a consensus based on other prediction tools such as MAPP, PolyPhen-1 and PolyPhen-2, SIFT, SNAP, and PANTHER. Therefore, a degree of accuracy is expected to be ensured. In addition, co-occurrence of mutations in NSP14 variants were also detected by the CoVal database.

#### 2.2.2 Prediction of RNA-interacting residues

Prediction of RNA-interacting residues in the SARS-CoV-2 NSP14 protein was made by the webserver Pprint. The webserver takes the amino acid sequence and automatically generates the evolutionary profile of whole sequence by running PSI-BLAST, generates support vector machine (SVM) pattern from this position-specific scoring matrix (PSSM) profile and then, predicts RNA interacting residues using SVM model[29]. The probability of correct prediction directly depends on the threshold, which by default set as -0.2. In this study, default threshold was used [29]. The red coloured residues are predicted as RNA-interacting and blue coloured residues are predicted as non RNA-interacting.

#### 2.2.3 Frequency distribution of amino acids and clustering

The frequency distribution of each amino acid present in a NSP14 sequence was determined using standard bioinformatics routine in *henson2004matlab*. For each NSP14 sequence, a twenty-dimensional frequency-vector considering the frequency of standard twenty amino acids can be obtained. The distance (Euclidean metric) between any two pairs of frequency vectors was calculated for each pair of NSP14 variants.

#### 2.2.4 Density-Based Clustering

Density-Based Spatial Clustering of Applications with Noise (DBSCAN) is one the most popular density-based clustering algorithm [30]. Twenty dimensional frequency vectors for all 3953 NSP14 sequences were considered as the dataset for clustering of 3953 SARS-CoV-2 NSP14 sequences. DBSCAN is a density-based clustering non-parametric algorithm: given a set of points in some space, it groups together points that are closely packed together. For the plotting purpose, we project the resulting clusters to a plane spanned by two principal components [30].

## 3. Results

### 3.1 Single Point Mutations in NSP14 Variants

Among the 527 amino acid residue positions, a total of 962 single mutations were identified within 3,953 unique NSP14 variants, as detailed in Tables (11 - 13). Notably, among these mutations, 398 were located within the ExoN domain, and 348 mutations were situated in the N7-MTase domain of the NSP14 sequence. Intriguingly, multiple mutations with frequencies ranging from 1 to 7 were observed at numerous residue positions, with seven distinct mutations documented at residues 132, 204, 211, and 496. A visual representation of mutation frequencies at each location is depicted in Figure 1.

**Figure 1.**
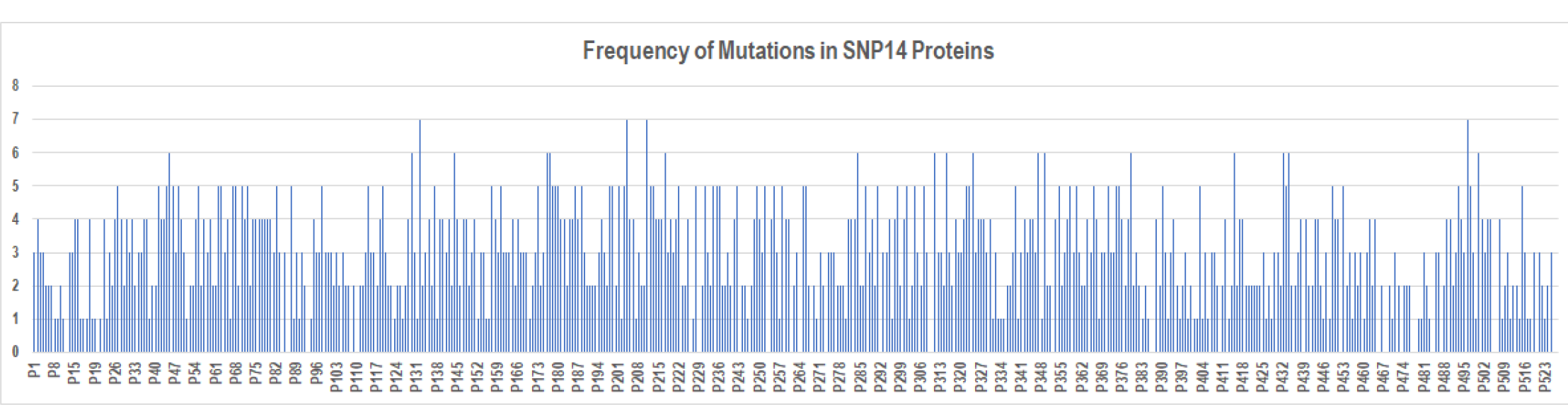
Total frequency of mutations per mutant residues of NSP14 reference sequence (YP 009725309)

Furthermore, a comprehensive analysis revealed a total of 414 deleterious mutations and 548 neutral mutations, classified based on pathogenicity, as outlined in Tables (11 - 13).

Notably, a variety of unique mutations were discovered across different geographic regions, as presented in Table 1. It’s worth mentioning that the highest number of unique single mutations within NSP14 variants were observed in the United States and the United Kingdom (Table 2). Additionally, it was observed that nearly half of the identified unique mutations were deleterious in the USA, UK, and India (Table 2).

**Table 1:** Unique single point mutations in NSP14 Variants in UK, USA, India, Canada, Switzerland, France, Spain, Mexico, Italy, Japan, and Denmark.

| UK |  |  | USA |  |  | India | Canada | Switzerland | France | Spain |
| --- | --- | --- | --- | --- | --- | --- | --- | --- | --- | --- |
| H427L | H148Q | L117P | I338L | F436I | A281G | A504Y | L177V | W227L | M241L | L411M |
| I294M | I242L | N306K | L406I | N129G | D331Y | D464G | Q145P | D30E | S407A | F377Y |
| D496H | I294S | Q313E | E204Q | N129Y | G59V | F367C |  | Y260F | T277A |  |
| G417S | K433N | S28N | K175R | D291G | K311T | H455S | Italy | Mexico | Japan | Denmark |
| R391S | R213S | V4A | V182A | D515A | N67K | S418V | W348R | S230T | A307P | Q513R |
| D324H | R525I |  | D179A | F89C | N71T | Y51C |  |  |  |  |
| E365G | Y361F |  | F350S | I332L | P451R |  |  |  |  |  |
| I166S | C382S |  | L322V | M57K | S369Y |  |  |  |  |  |
| C216L | I299M |  | N129A | S194C |  |  |  |  |  |  |

**Table 2:** Total number and percentage of deleterious and neutral mutations in different geo-locations.

| Unique mutations | UK | USA | India | Canada | Italy | Switzerland | Mexico | France | Japan | Spain | Denmark |
| --- | --- | --- | --- | --- | --- | --- | --- | --- | --- | --- | --- |
| Total mutations | 23 | 26 | 6 | 4 | 1 | 1 | 1 | 3 | 1 | 1 | 1 |
| Total Neutral mutations | 13 | 13 | 3 | 2 | 0 | 1 | 1 | 1 | 1 | 0 | 1 |
| Total Deleterious mutations | 10 | 13 | 3 | 2 | 1 | 0 | 0 | 2 | 0 | 1 | 0 |
| Neutral (%) | 56.52 | 50 | 50 | 50 | 0 | 100 | 100 | 33.3 | 100 | 0 | 100 |
| Deleterious (%) | 43.48 | 50 | 50 | 50 | 100 | 0 | 0 | 66.6 | 0 | 100 | 0 |

#### 3.1.1 Invariant Residues across NSP14 of various coronaviruses

Table 3 presents the invariant residues within the NSP14 reference sequence, emphasizing their consistency across 16 different coronaviruses originating from various hosts. The sequence homology analysis was conducted using Clustal-Omega, encompassing the following coronaviruses: Rousettus bat coronavirus HKU9 (YP_009924395.1), Betacoronavirus England (YP_009944304.1), Pipistrellus bat coronavirus HKU5 (YP_009944351.1), Tylonycteris bat coronavirus HKU4 (YP_009944322.1), SARS coronavirus Tor2 (NP_828871.1), Rabbit coronavirus HKU14 (YP_009924421.1), Murine hepatitis virus (YP_009915687.1), Rat coronavirus Parker (YP_009924380.1), Murine hepatitis virus (YP_009924354.1), Duck coronavirus (YP_009825026.1), Canada goose coronavirus (YP_009755922.1), Human coronavirus HKU1 (YP_460021.1), Murine hepatitis virus strain JHM (YP_209241.1), Bat coronavirus CDPHE15/USA/2006 (YP_008439224.1), Turkey coronavirus (YP_001941187.1), and the NSP14 reference sequence (YP_009725309.1).

**Table 3:** Invariant residues across NSP14 of various coronaviruses.

| Invariant Residue | Invariant Residue | Invariant Residue | Invariant Residue | Invariant Residue | Invariant Residue |
| --- | --- | --- | --- | --- | --- |
| 7 (L) | 112 (S) | 226 (C) | 280 (L) | 400 (R) | 452 (C) |
| 9 (K) | 114 (G) | 229 (H) | 281 (A) | 402 (D) | 466 (V) |
| 11 (C) | 118 (V) | 234 (D) | 286 (F) | 403 (T) | 468 (L) |
| 20 (P) | 123 (G) | 237 (Y) | 292 (W) | 404 (R) | 473 (C) |
| 34 (K) | 141 (P) | 238 (N) | 296 (Y) | 410 (N) | 474 (I) |
| 51 (Y) | 142 (P) | 239 (P) | 297 (P) | 411 (L) | 475 (T) |
| 56 (S) | 143 (G) | 243 (D) | 299 (I) | 413 (G) | 477 (C) |
| 59 (G) | 146 (F) | 245 (Q) | 302 (E) | 414 (C) | 478 (N) |
| 60 (F) | 148 (H) | 246 (Q) | 306 (N) | 416 (G) | 480 (G) |
| 73 (F) | 149 (L) | 247 (W) | 310 (R) | 417 (G) | 481 (G) |
| 75 (T) | 152 (L) | 248 (G) | 331 (D) | 418 (S) | 482 (A) |
| 79 (A) | 159 (W) | 251 (G) | 332 (I) | 419 (L) | 483 (V) |
| 83 (V) | 163 (R) | 253 (L) | 333 (G) | 420 (Y) | 484 (C) |
| 84 (R) | 166 (I) | 256 (N) | 334 (N) | 421 (V) | 487 (A) |
| 86 (W) | 169 (M) | 257 (H) | 335 (P) | 422 (N) | 488 (A) |
| 88 (G) | 172 (D) | 261 (C) | 336 (K) | 424 (H) | 491 (Y) |
| 89 (F) | 186 (W) | 264 (H) | 352 (D) | 425 (A) | 498 (Y) |
| 90 (D) | 191 (E) | 268 (H) | 355 (P) | 426 (F) | 499 (N) |
| 92 (E) | 192 (L) | 269 (V) | 368 (Y) | 427 (H) | 505 (G) |
| 95 (H) | 193 (T) | 270 (A) | 379 (D) | 428 (T) | 506 (F) |
| 102 (G) | 197 (Y) | 271 (S) | 380 (G) | 436 (F) | 509 (W) |
| 103 (T) | 198 (F) | 273 (D) | 388 (N) | 439 (L) | 514 (F) |
| 104 (N) | 200 (K) | 274 (A) | 389 (V) | 440 (K) | 518 (N) |
| 106 (P) | 202 (G) | 276 (M) | 392 (Y) | 443 (P) | 519 (L) |
| 108 (Q) | 207 (C) | 277 (T) | 393 (P) | 444 (F) | 520 (W) |
| 110 (G) | 210 (C) | 278 (R) | 398 (V) | 445 (F) |  |
| 111 (F) | 214 (A) | 279 (C) | 399 (C) | 447 (Y) |  |

These invariant residues signify regions of the NSP14 protein that exhibit a remarkable degree of conservation across diverse coronaviruses, regardless of their host origins.

A total of 160 invariant residue positions, identified through amino acid sequence homology comparisons with NSP14 sequences from various coronaviruses, were found in relation to the SARS-CoV-2 NSP14 reference sequence. Interestingly, it was observed that despite their invariance across other coronaviruses, several mutations were detected at most of these invariant residue positions within the different SARS-CoV-2 NSP14 sequences, as detailed in Table 4.

**Table 4:** Type of mutations detected in SARS-CoV-2 NSP14 variants at the invariant residues (Table-3)

| Invariant Residue | Type of mutation | Invariant Residue | Type of mutation | Invariant Residue | Type of mutation |
| --- | --- | --- | --- | --- | --- |
| 7 (L) | Deleterious | 214 (A) | Deleterious | 404 (R) | Deleterious |
| 34 (K) | Deleterious, Neutral | 234 (D) | Deleterious | 411 (L) | Deleterious |
| 51 (Y) | Neutral | 238 (N) | Deleterious | 413 (G) | Deleterious |
| 59 (G) | Deleterious | 239 (P) | Deleterious | 414 (C) | Deleterious |
| 75(T) | Deleterious | 245 (Q) | Deleterious | 416 (G) | Deleterious |
| 79 (A) | Deleterious | 247 (W) | Deleterious | 417 (G) | Neutral |
| 83 (V) | Deleterious | 248 (G) | Deleterious | 418 (S) | Deleterious |
| 88 (G) | Deleterious | 251 (G) | Deleterious | 420 (Y) | Deleterious |
| 89 (F) | Deleterious | 269 (V) | Deleterious | 421 (V) | Deleterious, Neutral |
| 95 (H) | Deleterious | 274 (A) | Deleterious | 424 (H) | Deleterious |
| 103 (T) | Deleterious | 276 (M) | Deleterious | 425 (A) | Deleterious |
| 111 (F) | Deleterious | 277 (T) | Deleterious | 427 (H) | Deleterious, Neutral |
| 112 (S) | Deleterious | 278 (R) | Neutral | 428 (T) | Deleterious |
| 114 (G) | Deleterious | 281 (A) | Deleterious | 436 (F) | Deleterious |
| 118 (V) | Deleterious, Neutral | 296 (Y) | Deleterious | 440 (K) | Neutral |
| 141 (P) | Deleterious | 297 (P) | Deleterious | 443 (P) | Deleterious |
| 142 (P) | Deleterious | 299 (I) | Deleterious | 444 (F) | Deleterious |
| 143 (G) | Deleterious | 306 (N) | Deleterious | 447 (Y) | Deleterious |
| 148 (H) | Deleterious | 331 (D) | Deleterious | 466 (V) | Neutral |
| 149 (L) | Deleterious | 332 (I) | Deleterious, Neutral | 474 (I) | Neutral |
| 152 (L) | Deleterious, Neutral | 336 (K) | Deleterious | 475 (T) | Deleterious |
| 159 (W) | Deleterious | 355 (P) | Deleterious | 481 (G) | Deleterious, Neutral |
| 163 (R) | Deleterious | 379 (D) | Deleterious, Neutral | 482 (A) | Deleterious |
| 166 (I) | Deleterious | 380 (G) | Deleterious | 488 (A) | Deleterious |
| 169 (M) | Deleterious | 388 (N) | Deleterious | 491 (Y) | Deleterious |
| 191 (E) | Deleterious | 389 (V) | Deleterious | 498 (Y) | Deleterious |
| 193 (T) | Neutral | 393 (P) | Deleterious | 506 (F) | Deleterious |
| 197 (Y) | Deleterious | 398 (V) | Deleterious | 509 (W) | Deleterious |
| 198 (F) | Deleterious | 399 (C) | Deleterious | 514 (F) | Deleterious |
| 200 (K) | Deleterious | 403 (T) | Deleterious |  |  |

Table 4 indeed reveals a noteworthy observation: nearly all of the invariant residues identified in Table 3, which were considered invariant across various coronaviruses, exhibited deleterious mutations within the SARS-CoV-2 NSP14 variants. This finding underscores the potential significance of these invariant positions in the functional and structural integrity of NSP14, as mutations in these positions are more likely to have detrimental effects on the virus.

#### 3.1.2 Invariant Residues across NSP14 Variants

In Table 5, we presented a list of 110 “Invariant residues” of the SARS-CoV-2 NSP14. These are amino acid residues that did not exhibit any single mutations in the unique NSP14 variants considered in this study, highlighting their remarkable stability and lack of variation within this specific dataset.

**Table 5:** 110 invariant residues positions in SARS-CoV-2 NSP14 variants (amino acid residues marked in bold font are invariant across all 16 NSP14 sequences of coronaviruses as found in Table 3)

| AA position of Invariant Residues in NSP14 Variants |  |  |  |  |  |  |  |  |
| --- | --- | --- | --- | --- | --- | --- | --- | --- |
| AA Position | AA Position | AA Position | AA Position | AA Position | AA Position | AA Position | AA Position | AA Position |
| 8 | **73** | **110** | **226** | **273** | 340 | **419** | 470 | **518** |
| **9** | **84** | **123** | **229** | **279** | 351 | **422** | **473** | **519** |
| **11** | **86** | 126 | **237** | **280** | **352** | **426** | **477** | **520** |
| 12 | **90** | 127 | **243** | 285 | 362 | **439** | **478** | 523 |
| 18 | **92** | **146** | **246** | **286** | **368** | **445** | 479 | 526 |
| 19 | 93 | 170 | **253** | **292** | 385 | 446 | **480** | 527 |
| **20** | 94 | **172** | **256** | **302** | 386 | **452** | **483** |  |
| 25 | **102** | **186** | **257** | 309 | 387 | 454 | **484** |  |
| 38 | **104** | **192** | **261** | **310** | **392** | 456 | **487** |  |
| **56** | 105 | **202** | **264** | **333** | **400** | 458 | 492 |  |
| **60** | **106** | **207** | **268** | **334** | 401 | 465 | **499** |  |
| 65 | **108** | 209 | **270** | **335** | **402** | 467 | **505** |  |
| 66 | 109 | **210** | **271** | 337 | **410** | **468** | 517 |  |

Based on the information presented in Table 5, it was deduced that 71 amino acid residues, marked in bold font, remained invariant across all 16 NSP14 sequences of various coronaviruses. In contrast, 37 amino acid residues (e.g., 8, 12, 18, 19, and others), which were not bolded in Table 5, were observed to be invariant in the unique 3953 SARS-CoV-2 NSP14 variants but exhibited variability across the other 16 different NSP14 coronavirus sequences, as detailed in Table 3.

Furthermore, the analysis of Table 5 led to the identification of double, triple, and quadruple invariant regions, as outlined in Table 6. It was noted that only two quadruple invariant regions, CNLG (amino acid positions: 477-480) and YNLW (amino acid positions: 517-520), contained both the PPNN ordered residues. This finding highlights the uniqueness and potential functional significance of these specific regions within the NSP14 protein.

**Table 6:** Invariant regions of double, triple, and quadruple sizes and their polarity (P and N denote polar and non-polar residues, respectively). Yellow and blue marked regions belong to the Exon and N7-MTase domains of SARS-CoV-2 NSP14.

| Double Residues | Polar-Non-Polarity | Double Residues | Polar-Non-Polarity | Triple Residues | Polar-Non-Polarity | Quadruple Residues | Polar-Non-Polarity |
| --- | --- | --- | --- | --- | --- | --- | --- |
| FK(8-9) | NP | CL(279-280) | PN | LHP(18-20) | NPN | CNLG(477-480) | PPNN |
| CS(11-12) | PP | CF(285-286) | PN | EGC(92-94) | PNP | YNLW(517-520) | PPNN |
| QV(65-66) | PN | CR(309-310) | PP | NLP(104-106) | PNN |  |  |
| DT(126-127) | PP | YD(351-352) | PP | QLG(108-110) | PNN |  |  |
| LC(209-210) | NP | FY(445-446) | NP | GNP(333-335) | NPN |  |  |
| NH(256-257) | PP | PL(467-468) | NN | WNC(385-387) | NPP |  |  |
| AS(270-271) | NP | VC(483-484) | NP | RFD(400-402) | PNP |  |  |
|  |  | LQ(526-527) | NP |  |  |  |  |

### 3.2 Co-occurring mutations in NSP14 variants

Beyond the single non-synonymous mutations, it was observed that a limited number of co-occurring mutations were present in NSP14 variants originating from various geographic locations. These co-occurring mutations suggest potential interactions or synergistic effects among specific amino acid residues within the NSP14 protein, warranting further investigation into their functional implications.

#### 3.2.1 Co-occurring mutations and their classification based on pathogenicity

Table 7 effectively tabulated the co-occurring mutations found in NSP14 variants from various geographic locations. These co-occurring mutations were distinguished as “unique co-occurring mutations” because they were exclusively identified with multiple frequencies (*≥*2) in their respective geographic locations. Notably, it was observed that all co-occurring mutations listed in Table 7 were also previously detected as single mutations, as indicated in Tables 10-12. Conversely, there were single mutations that had not yet been identified as co-occurring mutations, suggesting distinct mutation patterns and dynamics within the NSP14 variants.

**Table 7:**
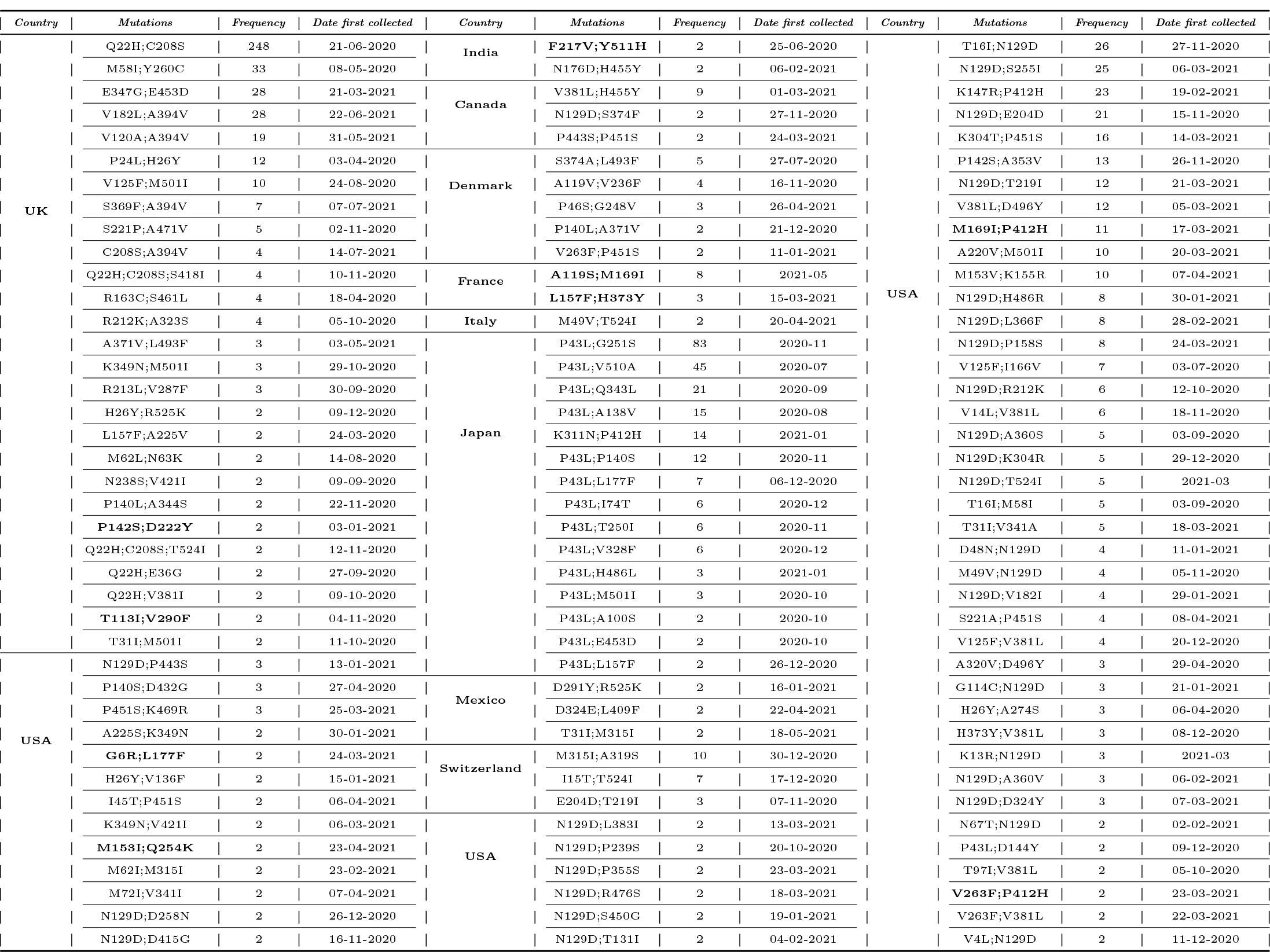
List of unique co-occurring mutations in different geo-locations. All the deleterious mutations occurring (such as G6R;L177F, H373Y;L157F and so on) together were marked bold.

Table 8 comprehensively presented the total number of co-mutations and unique co-mutations, along with their respective percentages, organized by geographic location as well as on a global scale. This tabulated data provides insights into the distribution and prevalence of these co-mutations across different regions, offering a broader perspective on the mutation landscape within NSP14 variants.

**Table 8:**
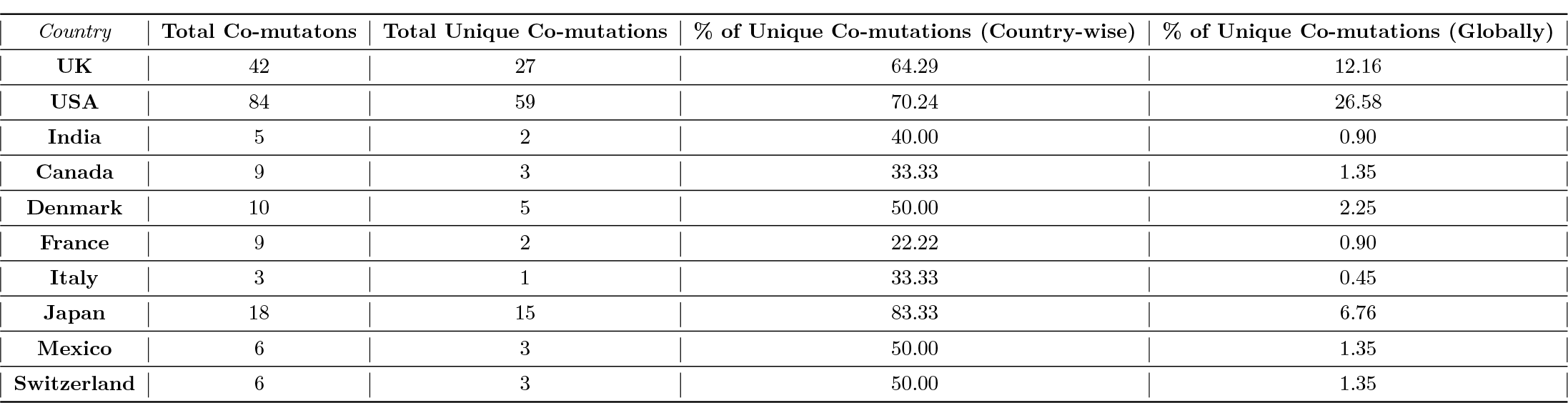
Frequencies and percentages of co-occurring mutations in NSP14 variants available in the various countries.

| Country | Total Co-mutations | Total Unique Co-mutations | % of Unique Co-mutations (Country-wise) | % of Unique Co-mutations (Globally) |
| --- | --- | --- | --- | --- |
| UK | 42 | 27 | 64.29 | 12.16 |
| USA | 84 | 59 | 70.24 | 26.58 |
| India | 5 | 2 | 40.00 | 0.90 |
| Canada | 9 | 3 | 33.33 | 1.35 |
| Denmark | 10 | 5 | 50.00 | 2.25 |
| France | 9 | 2 | 22.22 | 0.90 |
| Italy | 3 | 1 | 33.33 | 0.45 |
| Japan | 18 | 15 | 83.33 | 6.76 |
| Mexico | 6 | 3 | 50.00 | 1.35 |
| Switzerland | 6 | 3 | 50.00 | 1.35 |

The analysis of Table 8 revealed intriguing insights into unique co-occurring mutations across different geographic locations. Notably, the USA exhibited the highest percentage, with 26.58% of unique co-occurring mutations. The UK and Japan followed with 12.16% and 6.76%, respectively, while other geographic locations displayed significantly lower percentages of unique co-occurring mutations. Worth mentioning is the observation that out of the 18 detected co-occurring mutations in Japan, 15 were unique to that specific geographic location. Additionally, the USA had the next highest percentage of unique co-occurring mutations (Table 8).

To further elucidate the dynamics of co-occurring mutations within SARS-CoV-2 NSP14 variants, a co-occurring mutation flow diagram was created (Figure 2). This diagram provides a visual representation of how these mutations are interconnected. Additionally, the predicted pathogenicity of each mutation, whether neutral or deleterious, was indicated based on predictions from the PredictSNP web tool (Table 12).

**Figure 2.**
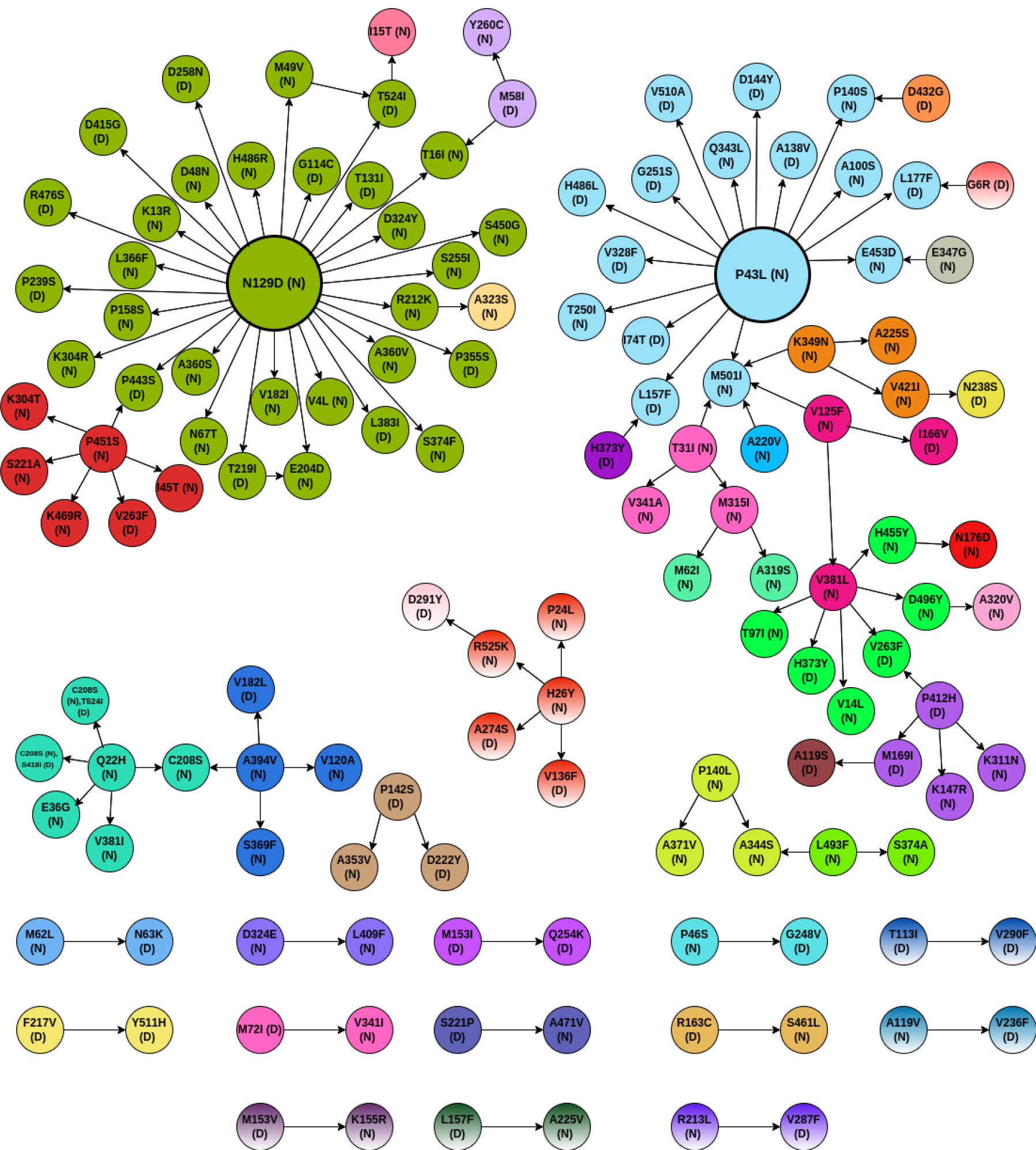
Co-mutation flow in SARS-CoV-2 NSP14 variants. The diagram represents the different co-mutations that are associated with each mutation. Note: 1. The colour coding has been done to understand which of the mutations are associated with a particular mutation. 2. The length of the arrow is just for representation and not indicative of anything. 3. For some mutations which are associated with more than one mutation, the colour coding of any one of the mutations is used. The **N** and **D** represent Neutral and Deleterious mutations, respectively.

Analysis of co-mutations in SARS-CoV-2 NSP14 variants revealed some interesting patterns. N129D emerged as the mutation with the highest number, with 31 co-mutations associated with it, followed by P43L with 15 co-mutations. Moreover, Q22H was identified with two triple-mutations, C208S and S418I, as well as C208S and T524I. Interestingly, both Q22H and C208S mutations were classified as neutral, while the additional co-occurring mutations, S418I and T524I, were deemed deleterious based on their pathogenicity.

Further classification of co-occurring mutations of length two was based on pathogenicity, resulting in three distinct classes: deleterious-deleterious (DD), deleterious-neutral (DN), and neutral-neutral (NN). Nine DD-type co-mutations, marked in bold in Table 7, were identified. Conversely, there were 47 DN-type and 61 NN-type co-mutations, revealing a range of mutation combinations with varying pathogenic implications (Table 9). These findings underscore the complexity of mutation interactions within NSP14 variants.

**Table 9:** Three different classes of co-occurring mutations in SARS-CoV-2 NSP14 variants.Yellow, blue, and green marked co-occurring mutations belong to the ExoN, N7-MTase, and both (one was from ExoN & another was from N7-MTase) domains, respectively.

| D-D | DN | DN | NN | NN | NN |
| --- | --- | --- | --- | --- | --- |
| M153I, Q254K | R213L, V287F | P43L, A138V | D324E, L409F | V125F, M501I | N129D, P158S |
| T113I, V290F | A394V, V182L | P43L, L177F | Q22H, C208S | M501I, K349N | N129D, K304R |
| F217V, Y511H | P142S, A353V | P140S, D432G | Q22H, V381I | K349N, A225S | N129D, A360S |
| P142S, D222Y | H26Y, V136F | N129D, T131I | Q22H, E36G | K349N, V421I | N129D, N67T |
| M169I, A119S | H26Y, A274S | N129D, T524I | A394V, C208S | M315I, A319S | N129D, V182I |
| P412H, M169I | P412H, K147R | T524I, I15T | A394V, V120A | M315I, M62I | N129D, E204D |
| P412H, V263F | P412H, K311N | N129D, D258N | A394V, S369F | T31I, M315I | N129D, V4L |
| L157F, H373Y | V263F, V381L | N129D, D415G | H26Y, R525K | T31I, V341A | N129D, S374F |
| L177F, G6R | V381L, H373Y | N129D, R476S | H26Y, P24L | M501I, A220V | N129D, A360V |
| DN | DN | DN | NN | NN | NN |
| N129D, G114C | V125F, I166V | N129D, P239S | P140L, A371V | M501I, T31I | N129D, R212K |
| M62L, N63K | V421I, N238S | N129D, P443S | P140L, A344S | P43L, M501I | R212K, A323S |
| P46S, G248V | P43L, L157F | N129D, T219I | L493F, A344S | P43L, T250I | N129D, S255I |
| A119V, V236F | P43L, I74T | T219I, E204D | L493F, S374A | P43L, Q343L | N129D, S450G |
| R163C, S461L | P43L, V328F | N129D, L383I | V381L, V14L | P43L, P140S | N129D, D324Y |
| A471V, S221P | P43L, H486L | N129D, P355S | V381L, D496Y | P43L, A100S | N129D, T16I |
| M72I, V341I | P43L, G251S | M58I, T16I | D496Y, A320V | P43L, E453D | P451S, K304T |
| M153V, K155R | P43L, V510A | M58I, Y260C | V381L, T97I | E453D, E347G | P451S, S221A |
| L157F, A225V | P43L, D144Y | M49V, T524I | V381L, H455Y | N129D, D48N | P451S, K469R |
|  | D291Y, R525K | P443S, P451S | H455Y, N176D | N129D, K13R | P451S, I45T |
|  |  | P451S, V263F | V125F, V381L | N129D, L366F | N129D, H486R |
|  |  |  |  |  | N129D, M49V |

**Table 10:**
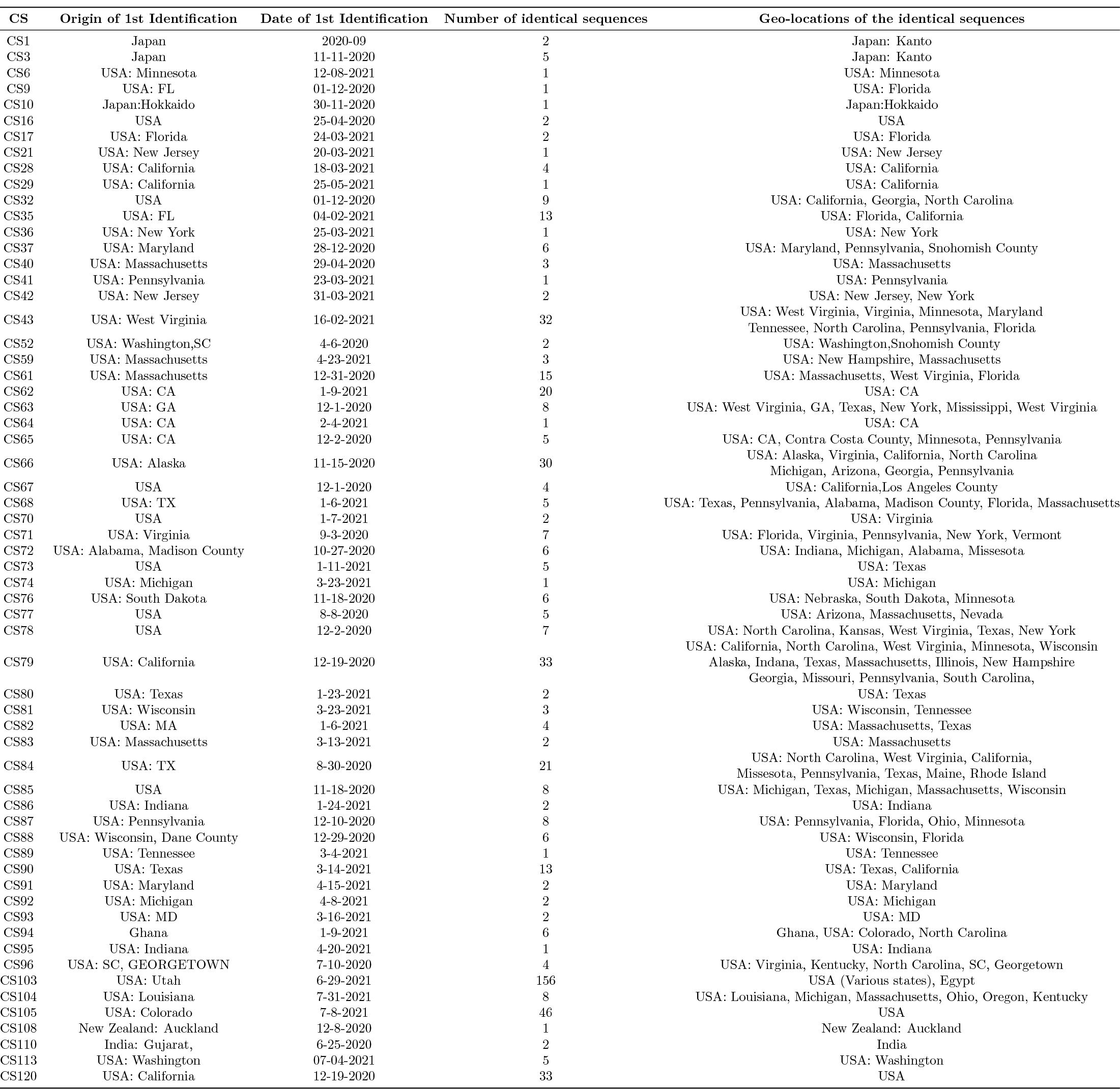
Geo-location and date of first identification and subsequent detection (with 100% identical) of co-mutated sequences.

The tracking of co-mutated sequences in SARS-CoV-2 NSP14 variants provides valuable insights into the spread and evolution of these mutations. In Table 10, the first detected geographic location and the respective dates of the co-mutated sequences are documented. Additionally, Table 10 presents the subsequent spread of these co-mutated sequences across other geographic locations. These details shed light on the geographical and temporal dynamics of co-mutations within NSP14 variants, contributing to the understanding of their emergence and dissemination.

#### 3.2.2 Phylogenetic variety of co-mutated NSP14 sequences

To gain a better understanding of the co-mutated sequences in Table 14, they were grouped into different clusters based on amino acid sequence homology with the reference SARS-CoV-2 NSP14 sequences. This clustering process is visualized in Figure 3, allowing for the identification of patterns and relationships among these sequences. Clustering based on sequence homology helps categorize co-mutated sequences with similar genetic characteristics, aiding in the analysis of their potential functional significance and evolutionary trends. 120 co-mutated sequences *CS*_1_ to *CS*_120_ were diversely allied into seven clusters. It turned out that reference SARS-CoV-2 NSP14 (YP_009725309) placed into seven distinct set of clusters as derived (and marked) in Figure 3 due to 120 co-occurring mutations. This diversity indicates wide span of varied functionalities of the NSP14 sequences.

**Figure 3.**
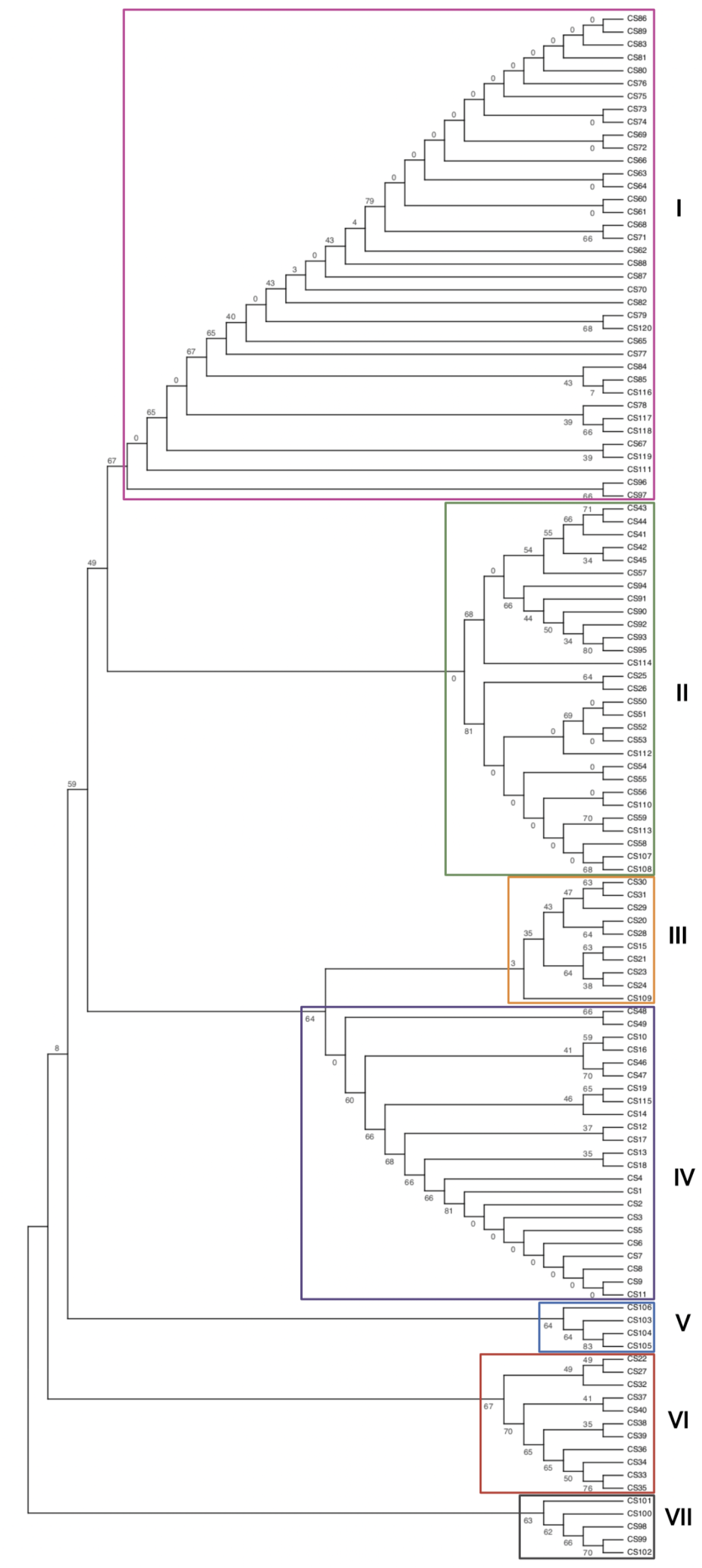
Phylogenetic relationship based on amino acid sequence homology of 120 co-mutated NSP14 sequences

### 3.3 Interacting residues of SARS-CoV-2 NSP14 sequences and mutations

A total of 146 interacting residues (marked red in Figure 4) among 527 residues present in the reference SARS-CoV-2 NSP14 sequence (YP_009725309) were predicted using the web-suite [29].

**Figure 4.**
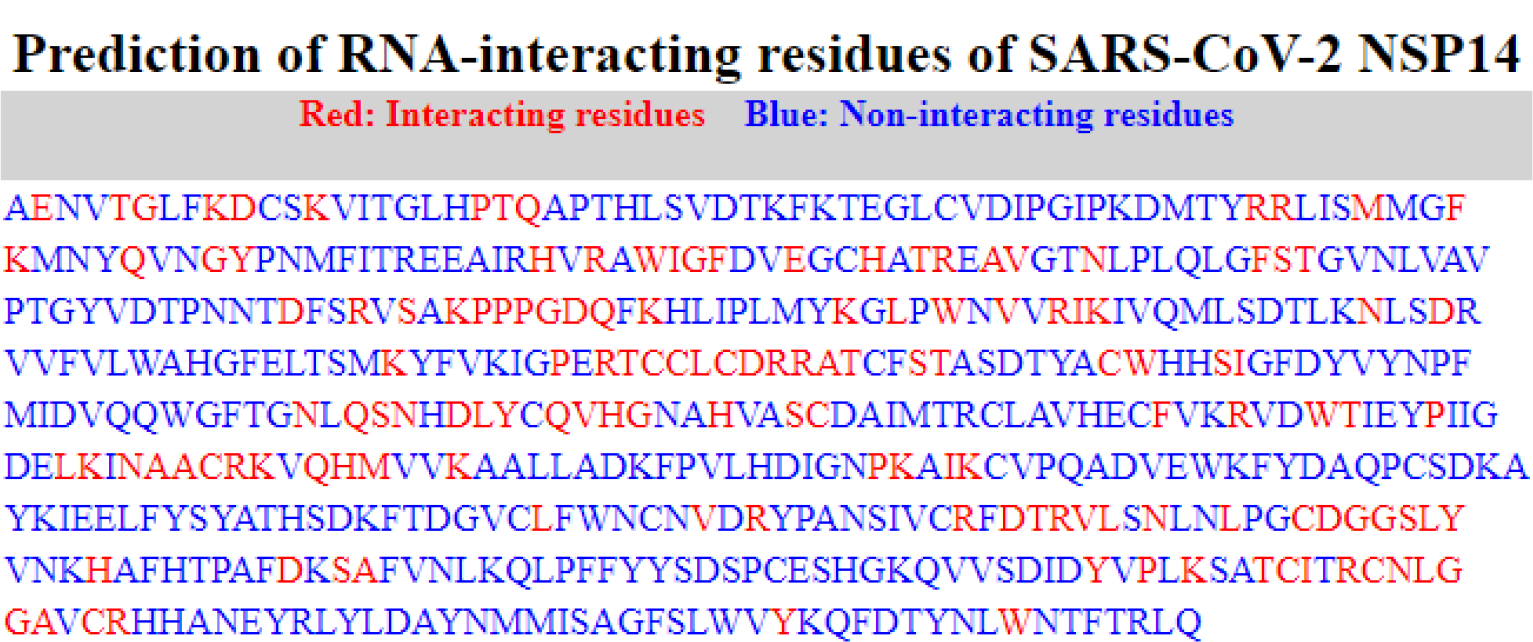
Interacting residues of wild-type SARS-CoV-2 NSP14 sequence

The analysis of the 146 interacting residues in Figure 4 revealed several interesting observations:

- Among these interacting residues, 113 possessed various mutations, indicating their potential role in the genetic diversity of NSP14.
- Specifically, 42 of these interacting residues were associated with only deleterious mutations, emphasizing their significance in terms of potential functional implications.
- On the other hand, 43 interacting residues had exclusively neutral mutations, suggesting that they may not directly impact the function or stability of the protein
- Additionally, a subset of 28 interacting residues exhibited mixed mutations, including both neutral and deleterious types, which could lead to complex effects on the the behaviour of the protein.
- Notably, a total of 33 interacting residues remained invariant, suggesting their crucial role in maintaining protein stability or function (Figure 5).

**Figure 5.**
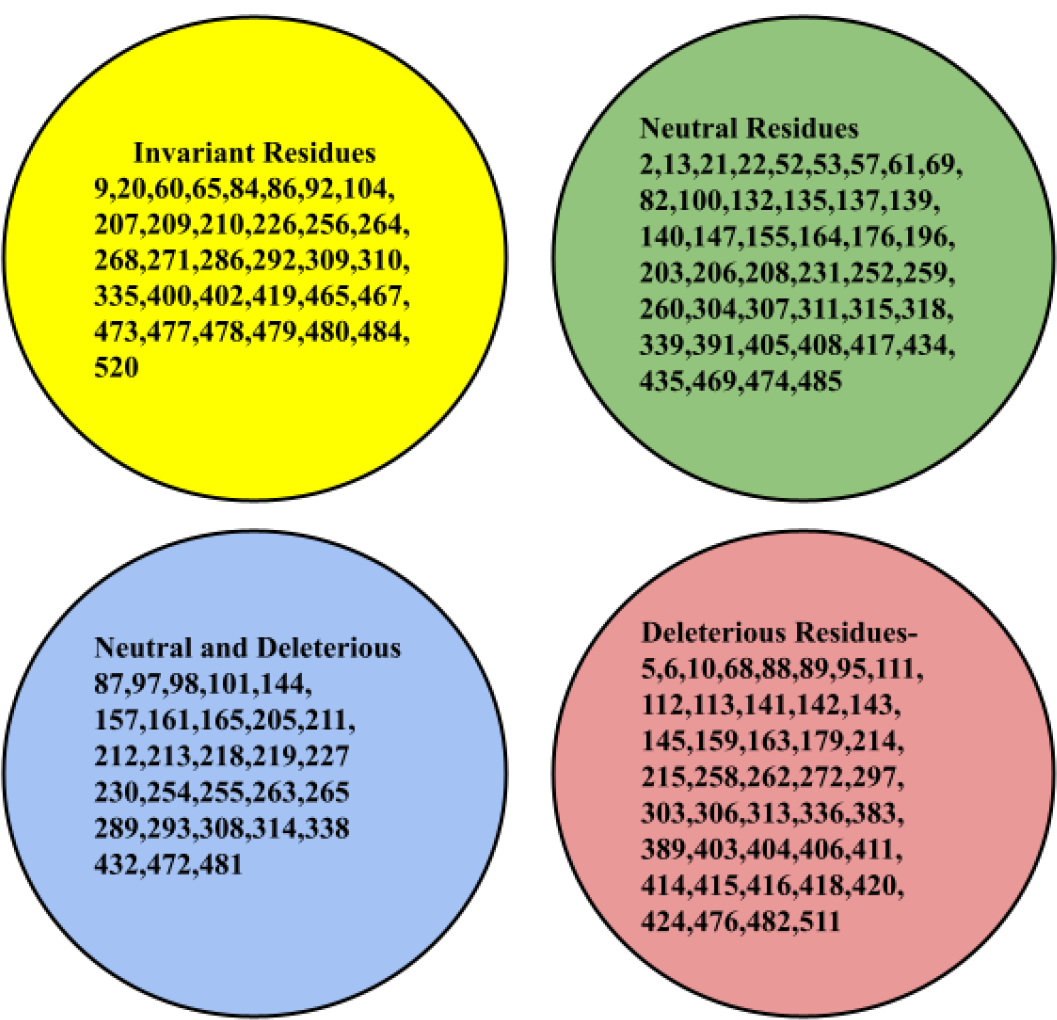
Three disjoint classes of interacting residues based on pathogenicity of mutations were detected. Invariant interacting residues set is marked in yellow.

These findings collectively provide insights into the interplay between mutations and interacting residues within NSP14, contributing to our understanding of its structural and functional aspects.

### 3.4. Variations and clustering of unique SARS-CoV-2 NSP14 variants

#### 3.4.1. Amino acid frequency distribution and clustering

To gain a deeper understanding of the quantitative variations among the 3953 unique SARS-CoV-2 NSP14 variants, several analyses were conducted:

A. Amino Acid Frequency Distribution: The frequency of each amino acid across all 3953 NSP14 sequences was computed and visualized, providing insights into the prevalence of specific amino acids within the variants (Figure 6(A)).
B. Pairwise Distance Calculation: Pairwise distances among all 3953 NSP14 variants were calculated, resulting in a distance matrix. This distance matrix was then plotted, revealing the genetic dissimilarity or relatedness among the variants (Figure 6(B)).

**Figure 6.**
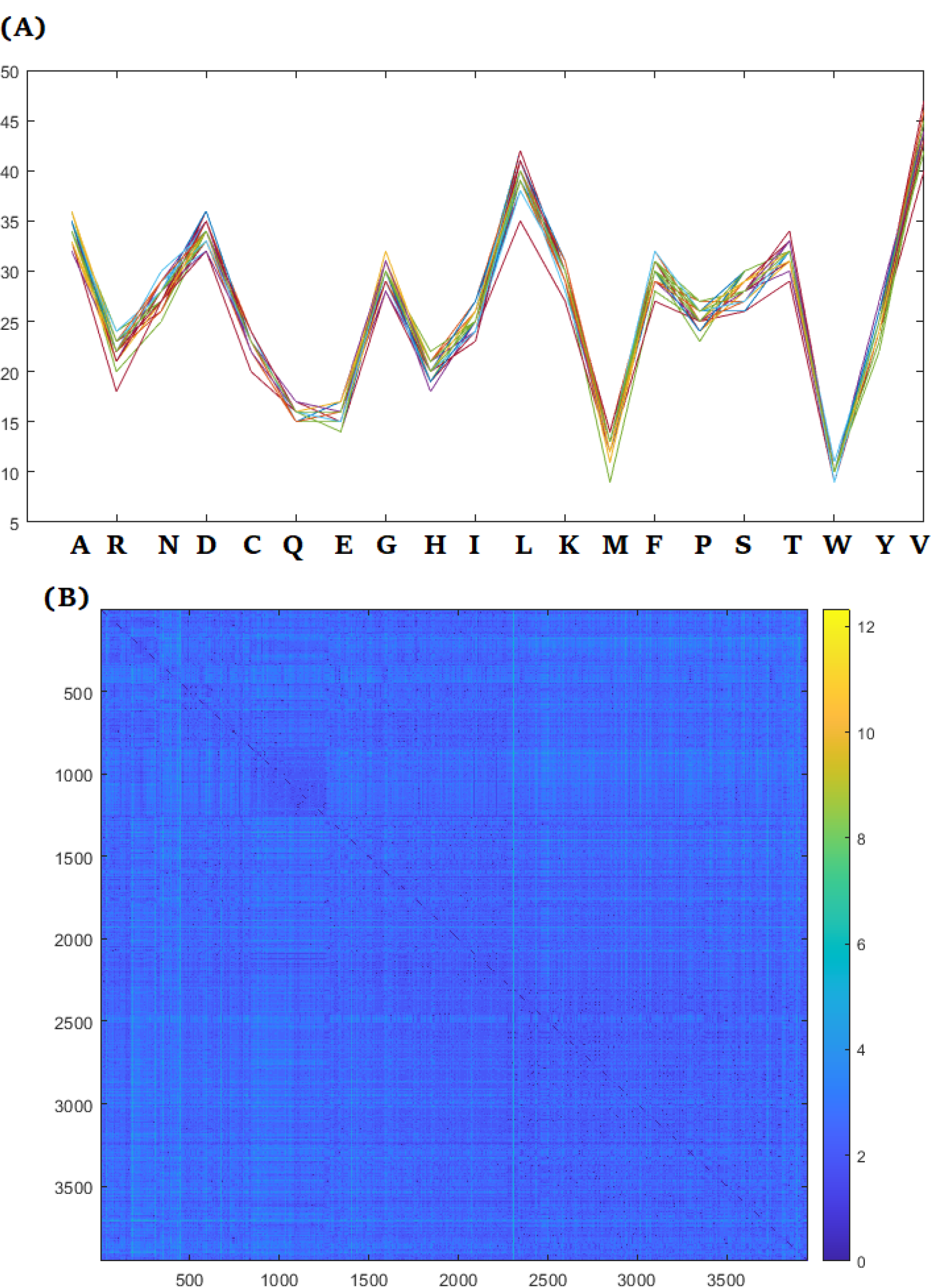
(A): Frequency distribution of each amino acids in 393 SARS-CoV-2 NSP14 sequences, (B): Pairwise distance among 3953 SARS-CoV-2 NSP14 sequences

The analysis of amino acid frequency distribution among the 3953 NSP14 sequences provided several interesting observations:

- Valine (V) was the most abundant amino acid, with a frequency ranging from 40 to 47 across the NSP14 variants.
- Conversely, tryptophan (W) was the least prevalent amino acid, with a presence ranging from 9 to 11 in the NSP14 variants.
- Amino acids Q and E had nearly identical frequencies, each occurring 16 times.
- Similar frequencies were also observed for the amino acid triplet (G, F, K) and the amino acid pairs (A, D), (R, C), and (P, Y).

Additionally, the clustering analysis revealed that the 3953 NSP14 sequences could be grouped into six distinct clusters: cluster:0, cluster:1, cluster:2, cluster:3, cluster:4, and cluster:5. These clusters contained varying numbers of NSP14 sequences, with cluster:0 being the largest, consisting of 2683 sequences, and cluster:5 being the smallest, with 16 sequences. A few sequences remained unclassified, denoted as cluster:-1 (Figure 7 and **Supplementary file-1**). These clusters provide insights into the genetic diversity and relatedness among the NSP14 variants.

**Figure 7.**
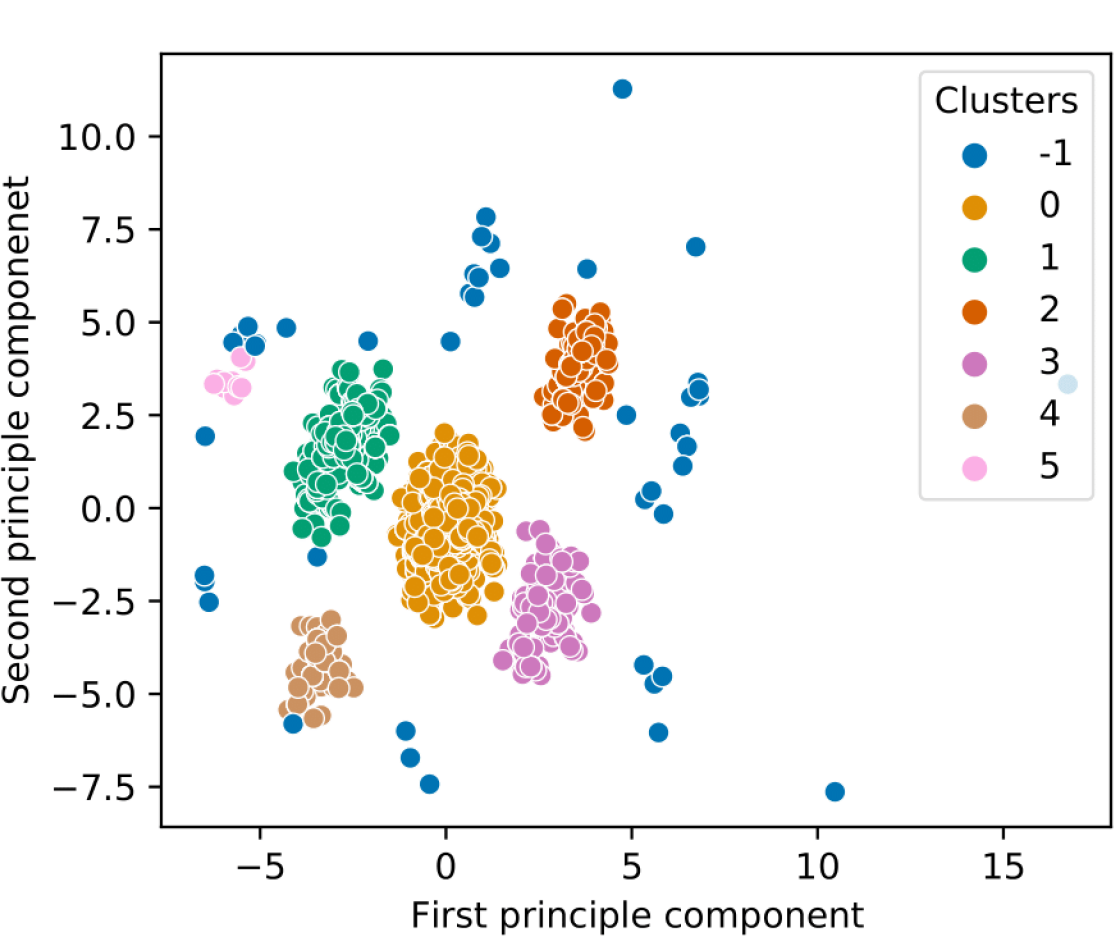
Clustering of NSP14 variants based on the amino acid frequency distribution of 3953 SARS-CoV-2 NSP14 sequences

The presence of 55 unclassified NSP14 sequences suggests that these sequences may not strongly align with any of the defined clusters. The existence of a dominant cluster indicates that the smaller clusters likely branched off recently from this dominant cluster, reflecting the evolutionary relationships among these variants.

It’s noteworthy that the unclassified cluster is equidistant from the remaining clusters, suggesting that these unclassified sequences may represent a distinct group with genetic characteristics that differ from those in the defined clusters. This equidistance indicates a unique evolutionary trajectory for this group of NSP14 sequences. Further analysis and investigation may provide insights into the specific genetic features and evolutionary history of these unclassified sequences.

#### 3.4.2. Quantitative physicochemical properties and clustering

The analysis of physicochemical properties for the 3953 SARS-CoV-2 NSP14 sequences provided valuable insights:

1. Isoelectric Points (PIs) Pattern: Among all the patterns of the measures plotted in Figure 8, the pattern of the isoelectric points (PIs) for the NSP14 variants exhibited a highly non-linear trend compared to the other quantitative measures obtained. This non-linearity in PIs suggests that the variations in charge distribution across the NSP14 sequences are distinct and potentially have functional implications.
2. Principal Component Analysis (PCA): Based on the quantitative features of each NSP14 sequence, a PCA analysis was conducted, resulting in the formation of three disjoint clusters (Figure 9). PCA is a powerful technique for dimensionality reduction and data visualization, allowing for the identification of underlying patterns and relationships within the dataset. These clusters provide a structured way to categorize and understand the variations in physicochemical properties among the NSP14 sequences.

**Figure 8.**
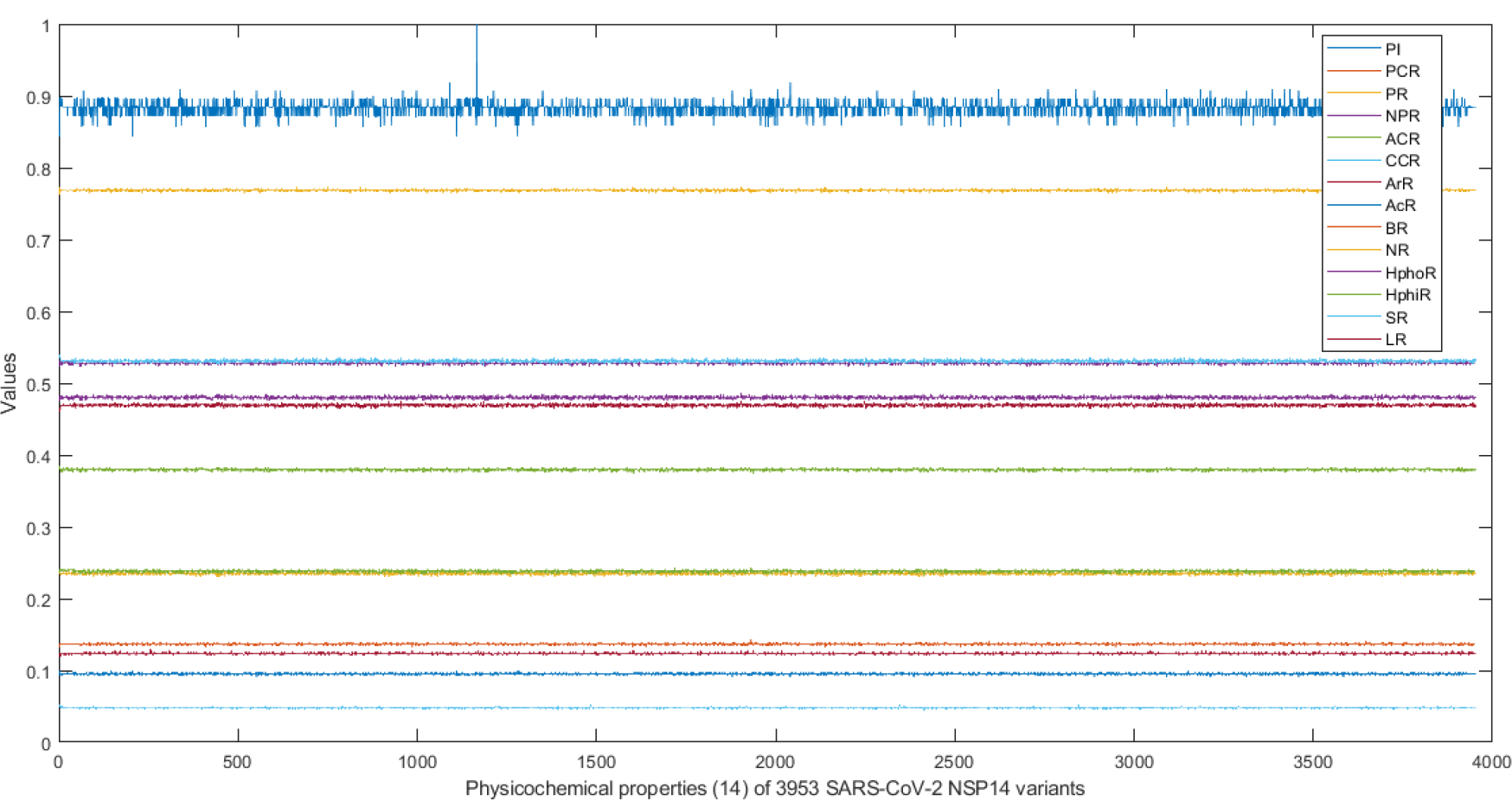
physicochemical features of 3953 SARS-CoV-2 NSP14 variants

**Figure 9.**
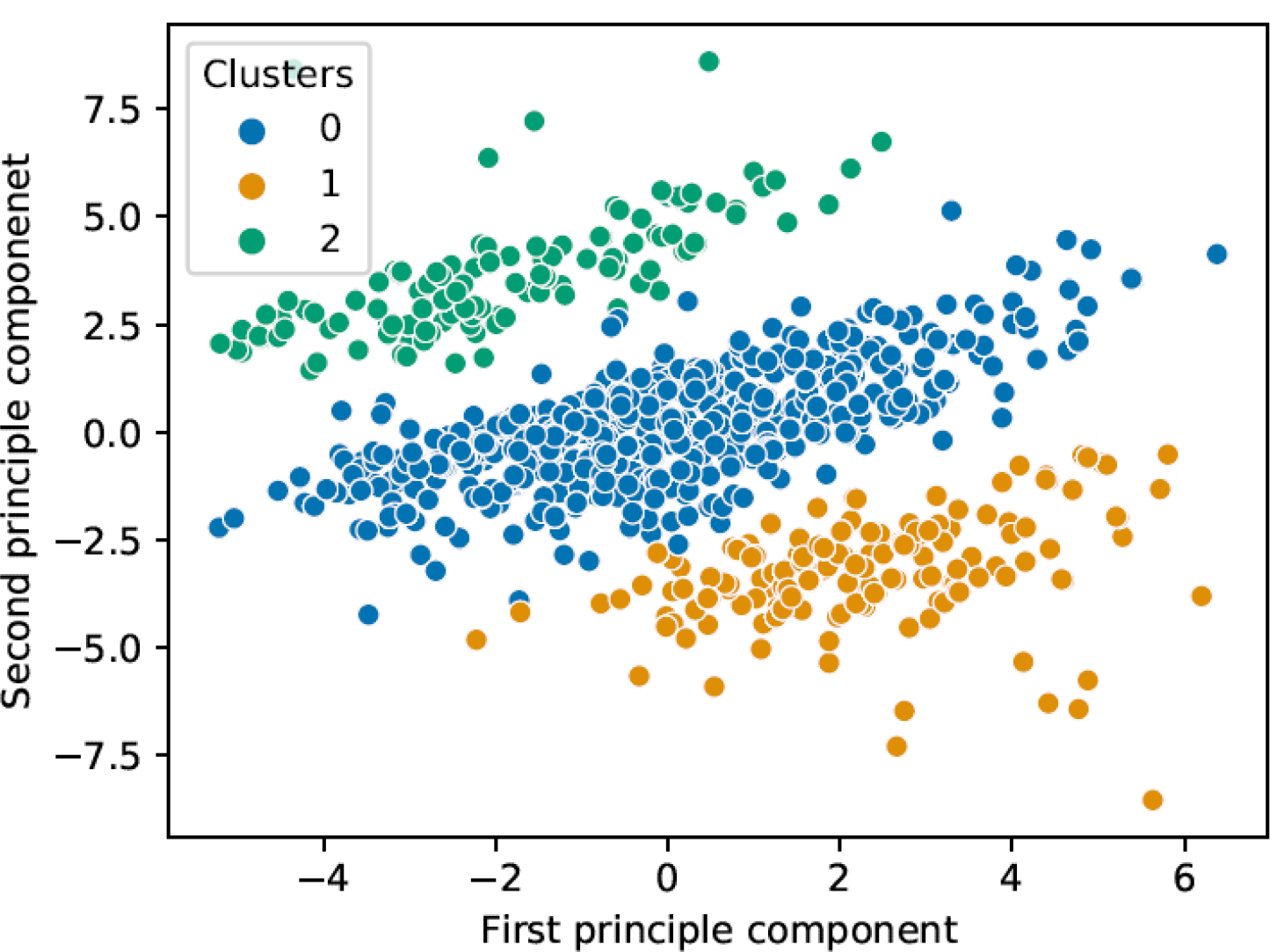
Clusters 0, 1, and 2 contain 3201, 297, and 455 respectively.

**Figure 10.**
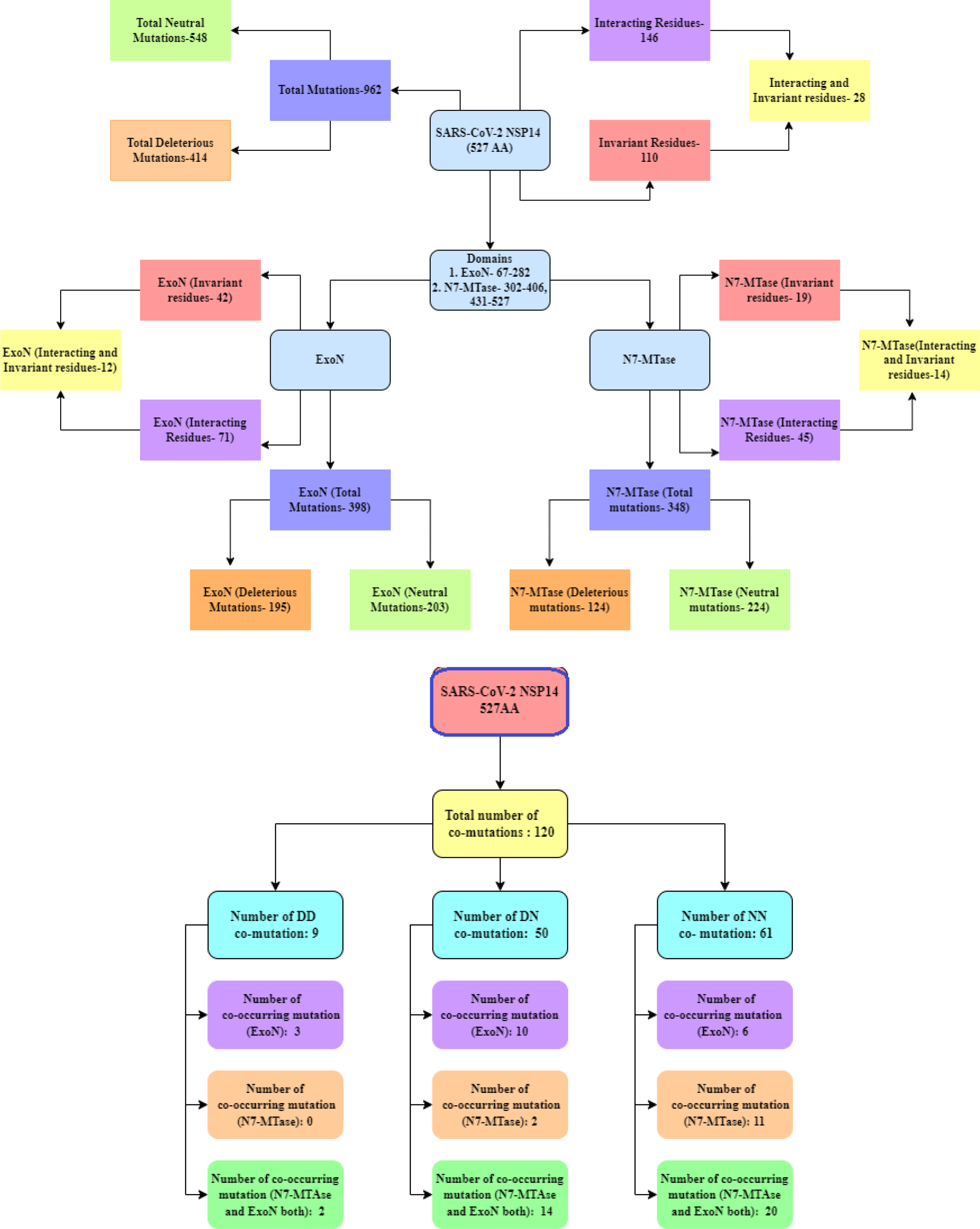
Schematic snapshot of frequency of mutations and co-occurring mutations, invariant residues, and interacting residues in ExoN and N7-MTase domains in SARS-CoV-2 NSP14

These analyses contribute to the understanding of the physicochemical characteristics and variations present within the NSP14 variants, shedding light on potential functional and structural implications.

In the PCA analysis, the majority of the NSP14 sequences, specifically 3201 out of 3953, were grouped into one cluster, referred to as cluster-0 (Supplementary file-3). Additionally, two smaller clusters, cluster-1 and cluster-2, were identified, containing 297 and 455 NSP14 variants, respectively. This clustering provides a structured way to categorize the NSP14 sequences based on their physicochemical properties, revealing distinct subgroups within the dataset. These subgroups may have unique characteristics and functional implications worth exploring further.

## 4. Discussion and Concluding Remarks

The NSP14 protein in SARS-CoV-2 is a nonstructural protein consisting of 527 amino acids and contains two distinct domains: the ExoN domain at the N-terminal and the N7-MTase domain at the C-terminal. The ExoN domain is particularly noteworthy for its proofreading activity, which plays a critical role in ensuring the fidelity of viral replication. NSP14 shares sequence similarity with other coronaviruses such as SARS-CoV and MERS-CoV [31].

Certain residues within NSP14 are typically conserved, but mutations in these residues can impact viral replication. Notably, mutations like R310A and F426A can affect viral replication, while H424A results in a crippled phenotype in SARS-CoV-2, MERS-CoV, and SARS-CoV [21]. The conservation of these residues, such as R310 and F426, across different NSP14 variants underscores their potential as drug targets, as mutations in these residues can influence viral phenotype and replication.. Analyzing the NSP14 sequences from various strains of SARS-CoV-2 revealed the presence of 962 mutations in total (Figure 4). Among these, 548 were classified as neutral mutations, and 414 were considered deleterious [32]. Deleterious mutations are those that result in a loss of gene function or impaired protein production, and they play a role in down-regulating the host immune response. It is worth noting that the balance between deleterious and neutral mutations in NSP14 is crucial for viral fitness and adaptation [33].

The ExoN domain of NSP14 exhibited a higher number of total mutations (398) compared to the N7-MTase domain (348 mutations). Inactivation of the ExoN domain can be lethal for SARS-CoV-2, and interestingly, ExoN mutations did not affect the activity of the N7-MTase domain [34]. Key residues in the ExoN region that interact with RNA are conserved across different coronaviruses, suggesting a shared RNA substrate recognition mechanism [35]. The N7-MTase domain of NSP14 is responsible for RNA capping and modulating host gene expression changes and translational repression [36, 23]. Mutations in key residues of the N7-MTase domain can interfere with viral replication, and these residues are conserved across different coronavirus families. Our analysis identified 19 invariant residues across the N7-MTase domain, with 14 of them being both invariant and interacting, indicating their crucial role in viral replication and RNA capping [21].

Co-mutations were also analyzed, and 120 co-mutations were identified across different strains. Neutral-neutral (NN) and deleterious-neutral (DN) co-mutations were the most common types. The presence of neutral mutations in comutation pairs can lead to stabilizing effects, promoting the survival and fitness of co-mutated strains [37]. Co-mutations can contribute to the development of different viral strains, increasing genomic diversity and potentially affecting viral fitness [38].

Overall, understanding the genetic variations and interactions within NSP14 is essential for unraveling the mechanisms underlying viral replication, evolution, and potential drug targets. Further research is needed to validate the impact of co-mutations on viral fitness and adaptation.

## Acknowledgements

We gratefully acknowledge the authors from the originating laboratories responsible for obtaining the specimens and the submitting laboratories where genetic sequence data were generated and shared via the NCBI and GISAID Initiatives, on which this research is based.

## Authors’ Contributions

SSH designed the study. SSH, DN, TB, PB contributed to the implementation of the research, to the analysis of the results. SSH, TB, and PB wrote the initial draft of the manuscript. All authors reviewed and edited. All authors read final version and approve.

## 5. Declaration of competing interest

Authors declare no competing interest to declare.

## Appendix

**Table 11:**
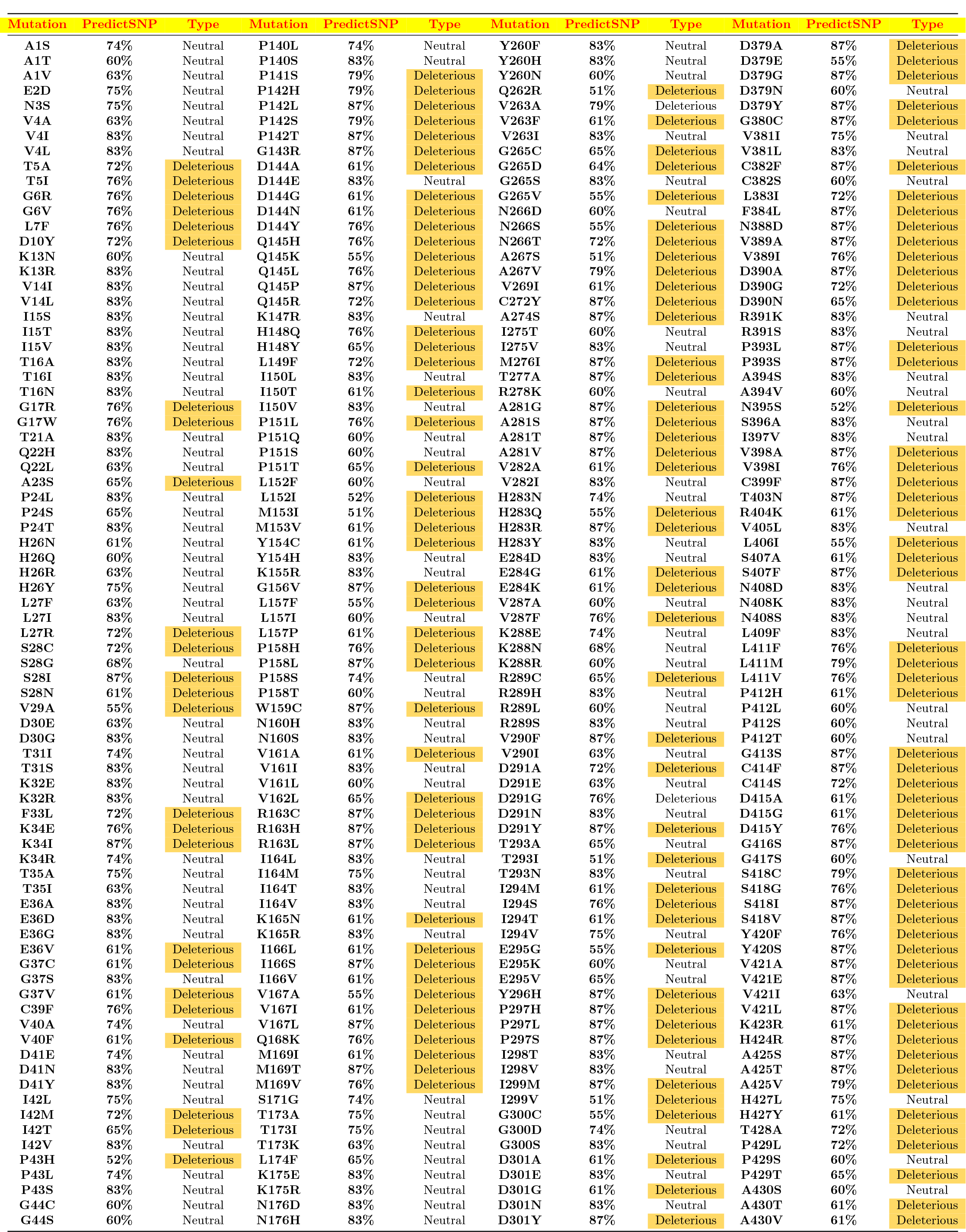
Pathogenicity (Predicted effect) of single mutations detected in the 3953 NSP14 variants.

**Table 12:** Pathogenicity (Predicted effect) of single mutations detected in the 3953 NSP14 variants (contd.)

| Mutation | PredictSNP | Type | Mutation | PredictSNP | Type | Mutation | PredictSNP | Type | Mutation | PredictSNP | Type |
| --- | --- | --- | --- | --- | --- | --- | --- | --- | --- | --- | --- |
| G44V | 83% | Neutral | N176S | 83% | Neutral | L303P | 55% | Deleterious | F431L | 74% | Neutral |
| I45L | 83% | Neutral | N176T | 74% | Neutral | K304M | 83% | Neutral | F431V | 76% | Deleterious |
| I45M | 60% | Neutral | L177F | 55% | Deleterious | K304N | 83% | Neutral | D432E | 60% | Neutral |
| I45T | 83% | Neutral | L177I | 83% | Neutral | K304R | 83% | Neutral | D432G | 72% | Deleterious |
| I45V | 83% | Neutral | L177V | 83% | Neutral | K304T | 75% | Neutral | D432N | 83% | Neutral |
| P46F | 62% | Deleterious | S178A | 63% | Neutral | I305L | 74% | Neutral | D432Y | 76% | Deleterious |
| P46L | 74% | Neutral | S178C | 76% | Deleterious | I305T | 76% | Deleterious | K433N | 61% | Deleterious |
| P46S | 83% | Neutral | D179A | 87% | Deleterious | I305V | 83% | Neutral | K433R | 83% | Neutral |
| K47N | 83% | Neutral | D179G | 87% | Deleterious | N306K | 87% | Deleterious | S434G | 83% | Neutral |
| K47R | 68% | Neutral | D179N | 87% | Deleterious | N306S | 76% | Deleterious | S434I | 83% | Neutral |
| D48G | 83% | Neutral | R180S | 83% | Neutral | A307P | 60% | Neutral | S434N | 83% | Neutral |
| D48N | 83% | Neutral | V181A | 76% | Deleterious | A307S | 83% | Neutral | A435S | 63% | Neutral |
| D48Y | 55% | Deleterious | V181I | 83% | Neutral | A307V | 65% | Neutral | A435V | 60% | Neutral |
| M49I | 83% | Neutral | V182A | 61% | Deleterious | A308S | 83% | Neutral | F436I | 87% | Deleterious |
| M49L | 74% | Neutral | V182I | 83% | Neutral | A308T | 65% | Deleterious | F436L | 65% | Deleterious |
| M49T | 74% | Neutral | V182L | 61% | Deleterious | A308V | 65% | Deleterious | V437A | 83% | Neutral |
| M49V | 83% | Neutral | F183L | 60% | Neutral | K311E | 74% | Neutral | V437F | 65% | Neutral |
| T50A | 60% | Neutral | V184F | 87% | Deleterious | K311N | 83% | Neutral | V437I | 83% | Neutral |
| T50I | 76% | Deleterious | V184I | 60% | Neutral | K311R | 83% | Neutral | N438S | 60% | Neutral |
| T50S | 83% | Neutral | L185S | 63% | Neutral | K311T | 83% | Neutral | K440R | 75% | Neutral |
| Y51C | 87% | Neutral | A187S | 63% | Neutral | V312F | 87% | Deleterious | Q441L | 83% | Neutral |
| R52K | 83% | Neutral | A187T | 87% | Deleterious | V312I | 60% | Neutral | L442S | 51% | Deleterious |
| R53K | 60% | Neutral | A187V | 87% | Deleterious | Q313E | 65% | Deleterious | L442V | 60% | Neutral |
| L54I | 55% | Deleterious | H188Q | 55% | Deleterious | H314P | 61% | Deleterious | P443L | 76% | Deleterious |
| I55T | 72% | Deleterious | H188R | 76% | Deleterious | H314Q | 74% | Neutral | P443S | 87% | Deleterious |
| I55V | 74% | Neutral | H188Y | 72% | Deleterious | H314Y | 60% | Neutral | P443T | 87% | Deleterious |
| M57I | 65% | Neutral | G189C | 76% | Deleterious | M315I | 74% | Neutral | F444Y | 87% | Deleterious |
| M57K | 74% | Neutral | F190L | 83% | Neutral | M315L | 83% | Neutral | Y447H | 87% | Deleterious |
| M57T | 75% | Neutral | E191D | 61% | Deleterious | M315T | 74% | Neutral | S448A | 61% | Deleterious |
| M57V | 65% | Neutral | T193A | 60% | Neutral | M315V | 83% | Neutral | D449E | 75% | Neutral |
| M58I | 72% | Deleterious | S194A | 63% | Neutral | V316F | 83% | Neutral | D449G | 76% | Deleterious |
| M58L | 74% | Neutral | S194C | 83% | Neutral | V316I | 83% | Neutral | D449N | 55% | Deleterious |
| G59C | 87% | Deleterious | M195I | 72% | Deleterious | V317A | 60% | Neutral | S450G | 83% | Neutral |
| G59S | 87% | Deleterious | M195T | 76% | Deleterious | V317I | 68% | Neutral | S450I | 79% | Deleterious |
| G59V | 87% | Deleterious | K196Q | 65% | Neutral | K318R | 83% | Neutral | S450N | 72% | Deleterious |
| K61R | 75% | Neutral | K196R | 83% | Neutral | A319S | 83% | Neutral | S450R | 51% | Deleterious |
| M62I | 83% | Neutral | Y197C | 87% | Deleterious | A319T | 87% | Deleterious | P451L | 63% | Neutral |
| M62L | 83% | Neutral | F198C | 87% | Deleterious | A319V | 51% | Deleterious | P451Q | 60% | Neutral |
| M62T | 52% | Deleterious | F198L | 87% | Deleterious | A320S | 72% | Deleterious | P451R | 72% | Deleterious |
| M62V | 74% | Neutral | V199A | 74% | Neutral | A320T | 72% | Deleterious | P451S | 68% | Neutral |
| N63K | 61% | Deleterious | V199L | 72% | Deleterious | A320V | 68% | Neutral | P451T | 68% | Neutral |
| N63S | 83% | Neutral | K200R | 55% | Deleterious | L321V | 74% | Neutral | E453A | 63% | Neutral |
| Y64C | 65% | Deleterious | I201L | 74% | Neutral | L322S | 74% | Neutral | E453D | 83% | Neutral |
| N67K | 74% | Neutral | I201M | 60% | Neutral | L322V | 75% | Neutral | E453G | 52% | Deleterious |
| N67S | 74% | Neutral | I201T | 76% | Deleterious | A323S | 83% | Neutral | H455Q | 83% | Neutral |
| N67T | 68% | Neutral | I201V | 74% | Neutral | A323V | 83% | Neutral | H455R | 83% | Neutral |
| G68C | 87% | Deleterious | P203H | 60% | Neutral | D324E | 71% | Neutral | H455S | 83% | Neutral |
| Y69H | 60% | Neutral | P203L | 74% | Neutral | D324G | 83% | Neutral | H455Y | 74% | Neutral |
| P70L | 63% | Neutral | P203S | 74% | Neutral | D324H | 65% | Neutral | K457R | 75% | Neutral |
| P70S | 83% | Neutral | P203T | 83% | Neutral | D324N | 83% | Neutral | V459I | 74% | Neutral |
| P70T | 74% | Neutral | E204A | 55% | Deleterious | D324Y | 83% | Neutral | V459L | 83% | Neutral |
| N71T | 83% | Neutral | E204D | 83% | Neutral | K325R | 83% | Neutral | V460L | 83% | Neutral |
| M72I | 51% | Deleterious | E204G | 72% | Deleterious | F326I | 83% | Neutral | S461A | 74% | Neutral |
| M72L | 83% | Neutral | E204Q | 74% | Neutral | F326L | 83% | Neutral | S461L | 60% | Neutral |
| M72V | 51% | Deleterious | E204V | 63% | Neutral | F326S | 83% | Neutral | D462E | 83% | Neutral |
| I74S | 87% | Deleterious | R205C | 61% | Deleterious | F326Y | 83% | Neutral | I463V | 83% | Neutral |
| I74T | 87% | Deleterious | R205H | 60% | Neutral | P327L | 74% | Neutral | D464G | 76% | Deleterious |
| I74V | 60% | Neutral | R205L | 83% | Neutral | P327Q | 83% | Neutral | D464N | 83% | Deleterious |
| T75A | 87% | Deleterious | T206I | 83% | Neutral | P327S | 83% | Neutral | D464Y | 76% | Neutral |
| T75I | 87% | Deleterious | T206S | 83% | Neutral | P327T | 83% | Neutral | V466L | 87% | Neutral |
| R76C | 87% | Deleterious | C208S | 83% | Neutral | V328F | 61% | Deleterious | K469R | 83% | Neutral |
| R76H | 87% | Deleterious | D211A | 65% | Neutral | V328I | 83% | Neutral | A471T | 74% | Neutral |
| R76L | 87% | Deleterious | D211G | 83% | Neutral | L329F | 55% | Deleterious | A471V | 60% | Neutral |
| R76S | 87% | Deleterious | D211Y | 65% | Deleterious | L329I | 83% | Neutral | T472A | 75% | Neutral |
| E77D | 83% | Neutral | R212G | 60% | Neutral | H330R | 72% | Deleterious | T472M | 65% | Deleterious |
| E77G | 51% | Deleterious | R212I | 61% | Deleterious | H330Y | 83% | Neutral | I474V | 71% | Neutral |
| E78A | 83% | Neutral | R212K | 83% | Neutral | D331Y | 87% | Deleterious | T475A | 79% | Deleterious |
| E78K | 60% | Neutral | R213C | 87% | Deleterious | I332L | 61% | Deleterious | R476C | 87% | Deleterious |
| A79S | 61% | Deleterious | R213H | 61% | Deleterious | I332V | 65% | Neutral | R476S | 87% | Deleterious |
| A79T | 87% | Deleterious | R213L | 60% | Neutral | K336R | 72% | Deleterious | G481C | 87% | Deleterious |
| I80K | 75% | Neutral | R213S | 74% | Neutral | I338L | 74% | Neutral | G481S | 63% | Neutral |
| I80T | 61% | Deleterious | A214T | 76% | Deleterious | I338T | 87% | Deleterious | G481V | 87% | Deleterious |
| I80V | 75% | Neutral | A214V | 76% | Deleterious | I338V | 75% | Neutral | A482V | 87% | Deleterious |
| R81K | 83% | Neutral | T215A | 61% | Deleterious | K339N | 60% | Neutral | R485G | 74% | Neutral |
| H82Q | 83% | Neutral | T215I | 61% | Deleterious | K339R | 83% | Neutral | R485K | 83% | Neutral |
| H82R | 83% | Neutral | C216F | 75% | Neutral | V341A | 83% | Neutral | H486L | 51% | Deleterious |
| H82Y | 83% | Neutral | C216L | 75% | Neutral | V341I | 83% | Neutral | H486R | 83% | Neutral |
| V83A | 61% | Deleterious | C216S | 61% | Deleterious | V341L | 60% | Neutral | H486Y | 65% | Deleterious |
| V83I | 76% | Deleterious | F217L | 51% | Deleterious | P342H | 83% | Neutral | A488D | 87% | Deleterious |

**Table 13:**
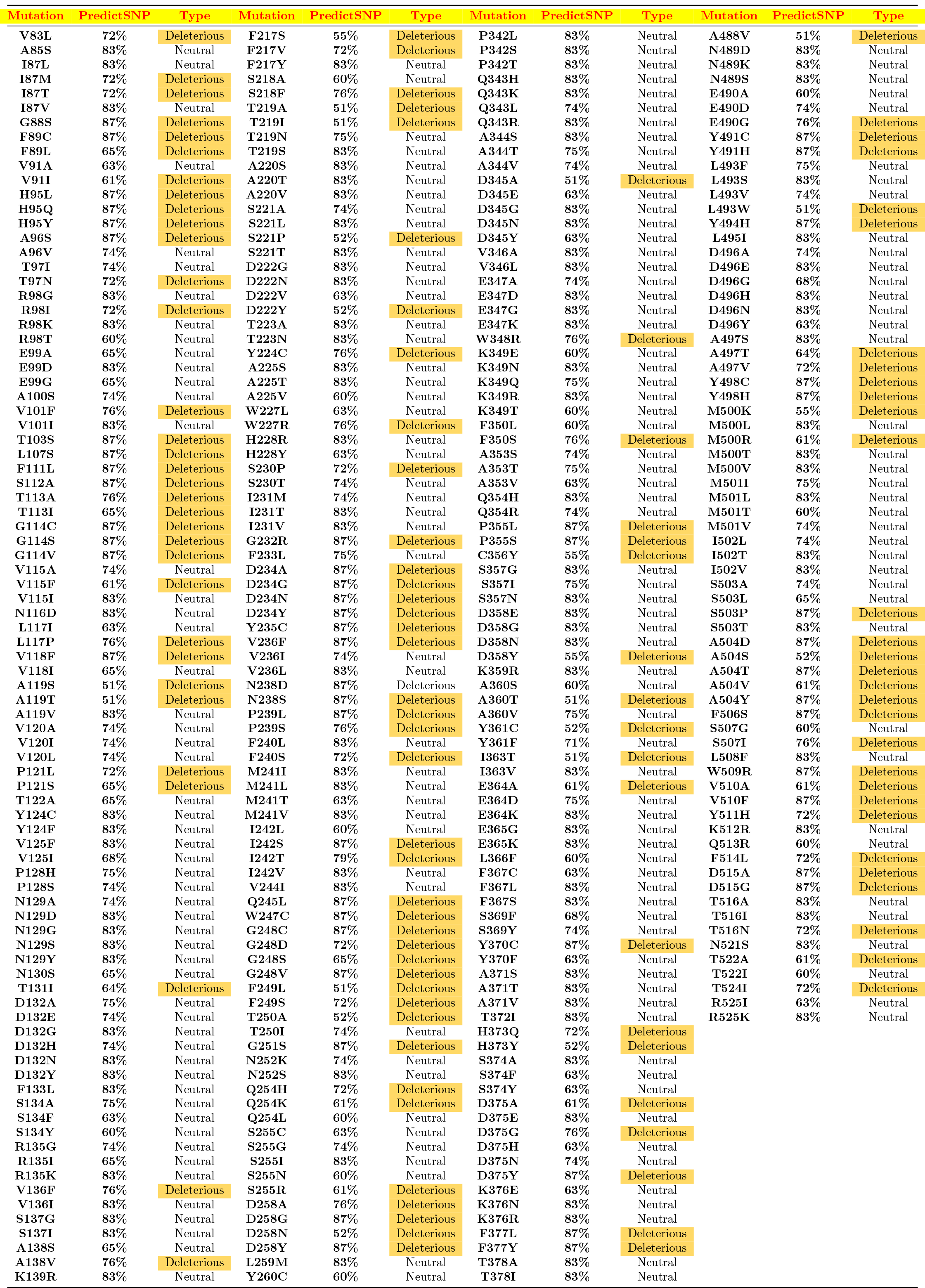
List of single point mutations and their pathogenicity.

**Table 14:**
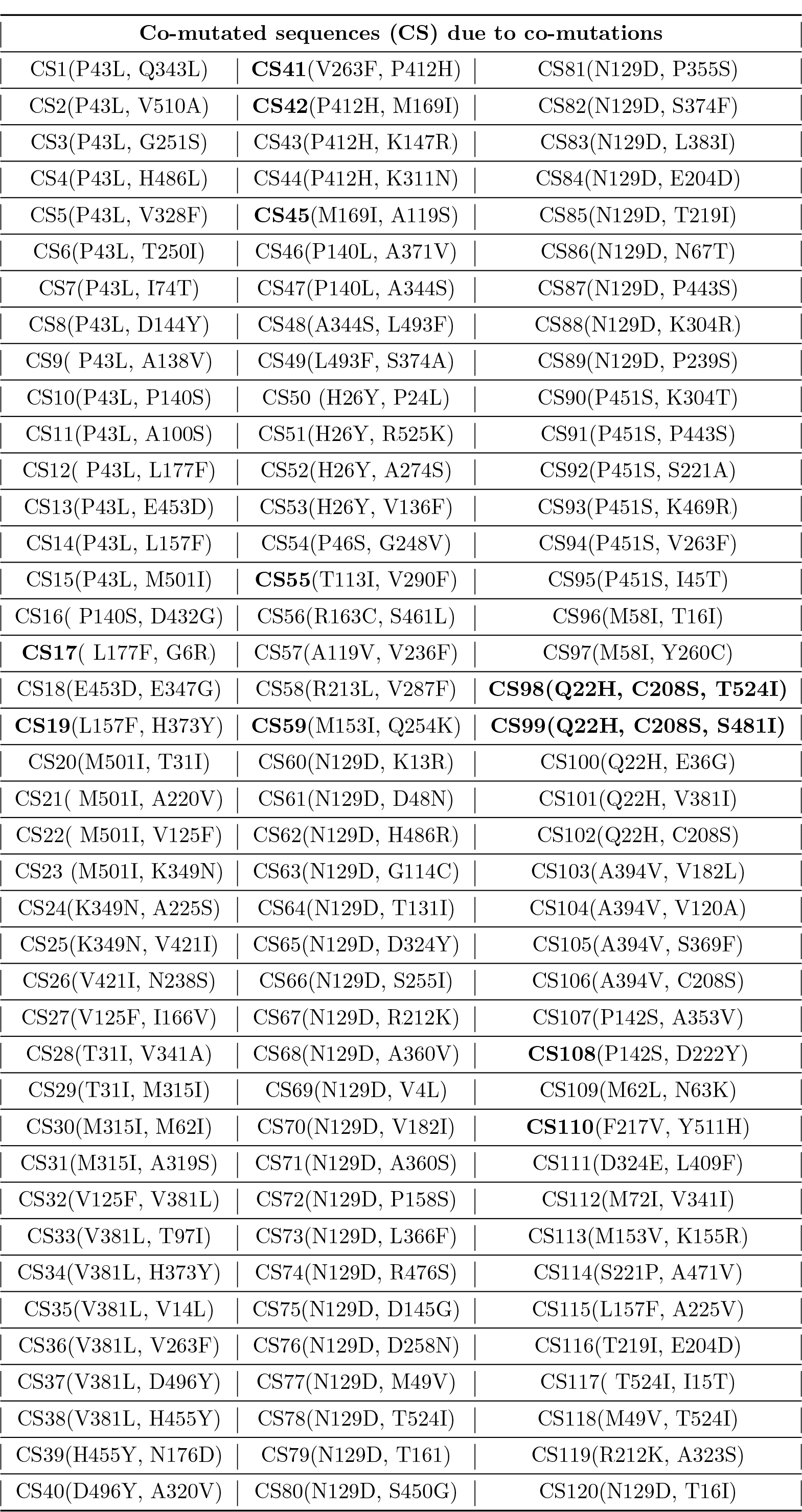
List of co-mutated sequences (CS) informatics. Nine bold marked CS were co-mutated sequences due to deleterious (DD type) mutations only.

| Co-mutated sequences (CS) due to co-mutations |  |  |
| --- | --- | --- |
| CS1(P43L, Q343L) | **CS41**(V263F, P412H) | CS81(N129D, P355S) |
| CS2(P43L, V510A) | **CS42**(P412H, M169I) | CS82(N129D, S374F) |
| CS3(P43L, G251S) | CS43(P412H, K147R) | CS83(N129D, L383I) |
| CS4(P43L, H486L) | CS44(P412H, K311N) | CS84(N129D, E204D) |
| CS5(P43L, V328F) | **CS45**(M169I, A119S) | CS85(N129D, T219I) |
| CS6(P43L, T250I) | CS46(P140L, A371V) | CS86(N129D, N67T) |
| CS7(P43L, I74T) | CS47(P140L, A344S) | CS87(N129D, P443S) |
| CS8(P43L, D144Y) | CS48(A344S, L493F) | CS88(N129D, K304R) |
| CS9( P43L, A138V) | CS49(L493F, S374A) | CS89(N129D, P239S) |
| CS10(P43L, P140S) | CS50 (H26Y, P24L) | CS90(P451S, K304T) |
| CS11(P43L, A100S) | CS51(H26Y, R525K) | CS91(P451S, P443S) |
| CS12( P43L, L177F) | CS52(H26Y, A274S) | CS92(P451S, S221A) |
| CS13(P43L, E453D) | CS53(H26Y, V136F) | CS93(P451S, K469R) |
| CS14(P43L, L157F) | CS54(P46S, G248V) | CS94(P451S, V263F) |
| CS15(P43L, M501I) | **CS55**(T113I, V290F) | CS95(P451S, I45T) |
| CS16( P140S, D432G) | CS56(R163C, S461L) | CS96(M58I, T16I) |
| **CS17**( L177F, G6R) | CS57(A119V, V236F) | CS97(M58I, Y260C) |
| CS18(E453D, E347G) | CS58(R213L, V287F) | **CS98(Q22H, C208S, T524I)** |
| **CS19**(L157F, H373Y) | **CS59**(M153I, Q254K) | **CS99(Q22H, C208S, S481I)** |
| CS20(M501I, T31I) | CS60(N129D, K13R) | CS100(Q22H, E36G) |
| CS21( M501I, A220V) | CS61(N129D, D48N) | CS101(Q22H, V381I) |
| CS22( M501I, V125F) | CS62(N129D, H486R) | CS102(Q22H, C208S) |
| CS23 (M501I, K349N) | CS63(N129D, G114C) | CS103(A394V, V182L) |
| CS24(K349N, A225S) | CS64(N129D, T131I) | CS104(A394V, V120A) |
| CS25(K349N, V421I) | CS65(N129D, D324Y) | CS105(A394V, S369F) |
| CS26(V421I, N238S) | CS66(N129D, S255I) | CS106(A394V, C208S) |
| CS27(V125F, I166V) | CS67(N129D, R212K) | CS107(P142S, A353V) |
| CS28(T31I, V341A) | CS68(N129D, A360V) | **CS108**(P142S, D222Y) |
| CS29(T31I, M315I) | CS69(N129D, V4L) | CS109(M62L, N63K) |
| CS30(M315I, M62I) | CS70(N129D, V182I) | **CS110**(F217V, Y511H) |
| CS31(M315I, A319S) | CS71(N129D, A360S) | CS111(D324E, L409F) |
| CS32(V125F, V381L) | CS72(N129D, P158S) | CS112(M72I, V341I) |
| CS33(V381L, T97I) | CS73(N129D, L366F) | CS113(M153V, K155R) |
| CS34(V381L, H373Y) | CS74(N129D, R476S) | CS114(S221P, A471V) |
| CS35(V381L, V14L) | CS75(N129D, D145G) | CS115(L157F, A225V) |
| CS36(V381L, V263F) | CS76(N129D, D258N) | CS116(T219I, E204D) |
| CS37(V381L, D496Y) | CS77(N129D, M49V) | CS117( T524I, I15T) |
| CS38(V381L, H455Y) | CS78(N129D, T524I) | CS118(M49V, T524I) |
| CS39(H455Y, N176D) | CS79(N129D, T161) | CS119(R212K, A323S) |
| CS40(D496Y, A320V) | CS80(N129D, S450G) | CS120(N129D, T16I) |

## Notes

### Competing Interest Statement

The authors have declared no competing interest.

